# Melanosomes degrade lipofuscin and precursors that are derived from photoreceptor membrane turnover in the retinal pigment epithelium—an explanation for the origin of the melanolipofuscin granule

**DOI:** 10.1101/2022.02.16.480523

**Authors:** Yanan Lyu, Alexander V. Tschulakow, Ulrich Schraermeyer

## Abstract

The accumulation of the age pigment lipofuscin within the retinal pigment epithelium (RPE) is one the most remarkable changes observed in association with age-related macular degeneration (AMD) and Stargardt disease. Both aging and pathological processes lead to the accumulation of melanolipofuscin (MLF) granules, which have been reported to reflect the onset of AMD more accurately than lipofuscin. The underlying mechanism by which MLF forms is still not understood. We investigate the potential role that melanin plays in the degradation of lipofuscin and MLF in pigmented *Abca4^-/-^* mice following treatment with several NO generating drugs. *Abca4^-/-^* mice are generally used as models for lipofuscin-related eye diseases. We also induced melanogenesis in albino *Abca4^-/-^* mice via the over-expression of tyrosinase, the key enzyme involved in melanogenesis. We compared the ultrastructure of lipofuscinogensis in the RPE of pigmented and albino *Abca4^-/-^* mice. Fluorescence microscopy was employed for the quantification of lipofuscin. We found high amounts of unique thin (3–4 nm) lamellar membranes (TLMs) that were left over from the degradation of photoreceptor disc membranes by high-resolution electron microscopy. Accumulated TLMs were significantly more frequent in the RPE cells of the albinos than the pigmented mice, indicating that melanin plays a role in removing TLMs. The intravitreal injection of several NO generating drugs was found to reduce the amount of autofluorescent lipofuscin in the cytoplasm of RPE cells, particularly the MLF granules of pigmented *Abca4^-/-^* mice. No effect was observed in terms of lipofuscin removal in NO-exposed albino *Abca4^-/-^* mice. However, transfection with tyrosinase led to a reduction in the lipofuscin levels of artificially pigmented RPE cells in albino *Abca4^-/-^* mice following the formation of melanin. The results show for the first time that melanin plays an important, if not a key, role in the degradation of lipofuscin in RPE cells.

## 1. Introduction

In this study, eyes from *Abca4^-/-^*^-^ mice, which are considered animal models for increased lipofuscinogenesis, Stargardt disease and dry AMD [1, 2] were examined by high-resolution electron microscopy.

Bisretinoids which are present in the RPE of these mice are lipofuscin precursors that are left over from the degradation of the photoreceptor membrane and are also related to the aging of the eyes. These molecules are formed during the metabolism of retinol in the outer segments of the photoreceptor under light exposure, and are better characterized than lipofuscin [3, 4]. Bisretinoids that are taken up by phagosomes into the RPE cannot be degraded by enzymes and are the origin of the lipofuscin that accumulates during aging. It seems that the lipofuscin load is regulated at a cellular level in normal aging eyes. High numbers of lipofuscin granules within the RPE have been associated with the development of RPE cell death and age-related macular degeneration. Both Stargardt patients and *Abca4^-/-^* mice have mutations in the ATP-binding cassette A4(Abca4) flippase that is located in the disc membranes of photoreceptors, which results in the gradual accumulation of high numbers of bisretinoids in the RPE. Nevertheless, the photoreceptors of such individuals work normally for a while, sometimes for decades, until the RPE becomes occupied by large amounts of lipofuscin, after which the RPE cannot work properly and visual acuity drops rapidly. Hence, unknown mechanisms may exist by which bisretinoids and lipofuscin are degraded. The ability to degrade lipofuscin and its precursors slows down progressively in aging humans and Stargardt mice models. This is exacerbated by the formation of granules composed of melanin and lipofuscin, or MLF granules, that also begin to accumulate and are particularly prominent in the RPE of aged human eyes, Stargardt patients, and Stargardt mice [1, 2, 5]. Feeny-Burns reported that MLF accumulation reflects the onset of AMD more closely than the accumulation of lipofuscin alone [6]. It has been hypothesized that MLF results from melanosomal autophagocytosis because it does not contain proteins derived from the photoreceptor membranes [7]. Although MLF granules seem to be important in the pathology of AMD, the origin of these granules has never been clarified. The melanosomes in the RPE belong to the lysosomal linage of cell organelles [8] and are known to possess many lysosomal enzymes [9]. Undegradable moieties that are associated with photoreceptor disc membranes are transported to the melanosomes and remain between the melanosomal membrane and the melanin core after residual materials originating from the disc membranes inside the melanosomes have been digested [10]. This process was demonstrated via the subretinal injection of gold-labeled rod outer segments [10], which showed that melanosomes have a role in lysosomal degradation. In this study we found high numbers of unique thin lamellar membranes (TLMs) that also resulted from incomplete degradation of the photoreceptor membrane in eyes from *Abca4^-/-^* mice. TLMs, which merit awareness because they may have pathological relevance, will be described in detail for the first time in this report.

This study hypothesizes that the indigestible bisretinoids follow the same pathway and can be degraded within the melanosomes. Studies have indicated that the lipofuscin component can be degraded by a variety of radicals or via the induction of chemical reactions during which radicals are formed, both in monkeys and *Abca4^-/-^* mice. Examples include the superoxide generators Soraprazan [11], Riboflavin [12], or light [13] and peroxidase [14]. The radical generator Visodyne, which was originally used to destroy pathological blood vessels in wet AMD, has also been found to efficiently remove the lipofuscin component from MLF granules in *Abca4^-/-^* mice following intravitreal injection [15]. Previous studies have indicated that the lipofuscin in the RPE can be degraded by radicals in vivo and that melanin can induce a variety of chemical reactions after coming in contact with nitric oxide (NO) [16]; we therefore investigated the effects that NO generating drugs have on the turnover of lipofuscin in *Abca4^-/-^* mice. To understand the role of melanin we compared the effects of NO generating drugs in albino and pigmented *Abca4^-/-^* mice. We present evidence for the involvement of melanosomes in the degradation of autofluorescent lipofuscin.

## 2. Methods

### 2.1. Reagents

3-morpholinosydnonimine(SIN-1) was purchased from Focus Biomolecules(Plymouth Meeting, PA USA). It could spontaneously release nitric oxide(NO) and superoxide anion[17]. Isosorbide dinitrate (ISDN), was purchased from Merus Labs Luxco II S.A R.L.(Grand Duchy Of Luxembourg). It is a medication used for angina pectoris by its ability to convert to nitric oxide[18]. Horseradish peroxidase was obtained from Sigma-Aldrich (Darmstadt, Deutschland).

### 2.2. Animals

Pigmented *Abca4^-/-^* mice (129S4/SvJae-Abca4^tm1Ght^) and albino *Abca4^-/-^* mice (BALB/cAbca4tm1Ght) were used in this work. Pigmented *Abca4^-/-^* mice (129S4/SvJae-Abca4^tm1Ght^) were purchased from Charles River (Sulzfeld, Germany). Albino *Abca4^-/-^* mice (BALB/c-Abca4 tm1Ght) were kindly donated by R. Radu (University of California, Los Angeles, CA). All strains were bred in our in-house facility in a 12h lights (approximately 50 lx in cages)/12h dark cycle with food and water ad libitum. All procedures were conducted with the approval of the local agency for animal welfare (Einrichtung für Tierschutz, Tierärztlichen Dienst und Labortierkunde der Eberhard Karls Universität Tübingen, Tuebingen, Germany) and local authorities (Regierungspräsidium Tübingen, Tuebingen, Germany) under the authorization reference number AK1/16.

### 2.3. Generation of Ad-Tyr

The adenoviral vector Ad-Tyr carries the inserted human tyrosinase cDNA. It was generated according to the published method [19].

### 2.4. Subretinal injection with Ad-Tyr

Twelve 3-4 month-old albino *Abca4^-/-^* mice were used for the experiments. The mice were anesthetized using isoflurane (Isoflurane CP®, CP-Pharma, Germany) inhalation (3.5% isoflurane and 25% oxygen). Pupils were dilated with tropicamide drops (Pharmacy of the University of Tuebingen, Germany) and a drop of topical anesthetic Novesine (OmniVision, Puchheim, Germany) was applied. Methocel 2% (OmniVision, Puchheim, Germany) was used to avoid drying corneas. The mice were positioned under surgical microscopy equipped with illumination. The eyeball position was adjusted by holding the conjunctiva with tweezers and gently pulling. The subretinal injection was performed by inserting the tip of the syringe tangentially into the eyes through the sclera into the subretinal space without damaging the lens or posterior retina. A volume of 1.5µl of Ad-Tyr (10^8^ iu/ µl) was injected through a 10 μL NanoFil syringe with NanoFil 34-gauge blunt needle (World Precision Instruments) into the subretinal space of the right eye of the mice.

The left eye of each animal remained untreated. Antibiotic ointment (GentamicinPOS®, Ursapharm, Saarbruecken, Germany) was applied routinely after injection. Two weeks after injection the fundus autofluorescence images were acquired in vivo. Afterwards, the mice were sacrificed and the eyes from all mice were enucleated and prepared for histology.

### 2.5. Intravitreal injection

Twenty 6 to 24 month-old mice in total were used in the experiments. The right eyes of mice were injected intravitreally with SIN-1 (65μg/μL) dissolved in PBS, the left eye of each animal remained untreated. Preoperative preparations were carried out as mentioned above. A small opening was made into the sclera at the pars plana region of the eyes by the tip of the syringe. A volume of 2 μL SIN-1 was injected intravitreally through the incision using a 10 μL NanoFil syringe with NanoFil 34-gauge beveled needle. To avoid reflux out of the injection site, the needle was retrieved very steadily. The eyeball was brought back into its normal position, and the antibiotic ointment was applied to the eye. Animals were kept for 2 days before sacrifice. Horseradish peroxidase and ISDN treatments were performed in the same manner. (Horseradish peroxidase 70ug/uL dissolved in PBS, ISDN 40μg/ uL dissolved in PBS).

### 2.6. Fundus autofluorescence image acquisition

Before the sacrifice of the Ad-Tyr treated mice, confocal scanning laser ophthalmoscopy (cSLO; Spectralis HRA+OCT, Heidelberg Engineering, Heidelberg, Germany) was used for fundus AF image acquisition as described previously[5]. Briefly, the mice were anesthetized intraperitoneally with narcosis (0.05 mg/kg of fentanyl, 5 mg/kg of midazolam, and 0.5 mg/kg of medetomidine). Methocel was applied to the cornea after fully dilated pupils. Detailed settings were as reported before. A 78-DPT non-contact slit lamp lens (Volk Optical, Inc., Mentor, OH 44060, USA) was fixed directly in front of the device. Moreover, a custom-made contact lens (100 DPT) was used on the cornea. The mouse was placed on an adjustable three-dimensional platform.

To align the camera and find the areas of interest, the near-infrared reflectance (NIR-R) mode was performed first. Short-wavelength autofluorescence (SW-AF) images and near-infrared autofluorescence (NIR-AF) were recorded at the same time according to previous settings.

### 2.7. Sample preparation for fluorescence and electron microscopy

Animals were sacrificed by isoflurane inhalation and subsequent cervical dislocation and the eyes were instantly enucleated, followed by fixation in 5% glutaraldehyde overnight at 4°C. To prepare for embedding, the eyecups were cut vertically into two halves through the optic disc after removing the cornea and lens. The eyes are embedded in Epon resin and later sectioned according to standard procedures[1, 20]. Reagents were purchased from AppliChem (Darmstadt, Germany), Serva (Heidelberg, Germany) and Merck (Darmstadt, Germany). For electron microscopy and light microscopy, the half was postfixed with 1% OsO4 in 0.1 M cacodylate buffer (pH 7.4) and stained with UAC. Ultra-thin sections (70nm) were stained with lead citrate and examined by electron microscope (Zeiss 900, Jena, Germany). For fluorescence analysis, post-fixation and heavy metals staining were omitted. Semi-thin sections (500nm) were prepared and cover-slipped with Dako fluorescent mounting medium then investigated by fluorescence microscopy.

### 2.8. Electron microscopic investigation and quantification of vacuoles containing TLMs

Sections were examined comprehensively for changes in RPE and choroid. The area of vacuole-like structures(described by Taubitz et al.)[1] and the length of the RPE layer were measured in 7-month-old pigmented and albino *Abca4^-/-^* mice. Thin lamellar membranes were measured in electron micrographs at 140000-fold magnification. All measurements were performed with imageSP Software (Minsk, Belarus)

### 2.9. Quantification of the area in RPE cytoplasm occupied by lipofuscin-like material

The images of ultrathin sections through the optic nerve head were recorded using electron microscopy for quantification of lipofuscin-like material. Ten electron micrographs of the RPE (magnification× 7.000) per eye were taken starting from the optic nerve head. The areas of lipofuscin-like material (derived from both lipofuscin and melanolipofuscin granules; for the melanolipofuscin, only the lipofuscin part of the granules was measured) were quantified in 6-24 months SIN-1 treated pigmented and albino *Abca4^-/-^* mice in an average of 5 eyes per group. Fiji software was used for the measurements. When the cytoplasm areas were measured, nuclei, microvilli, and basal labyrinth were excluded.

### 2.10. Fluorescence microscopy

Sections were examined using Zeiss Axioplan 2 microscope (Zeiss, Jena, Germany) equipped with a Lumencor Sola SE II NIR (Beaverton, OR, USA) light source. Filter setting details were as described previously[2]. To allow comparison of fluorescence intensities, acquisition times, as well as microscope and software settings were held constant for any set of samples. For fluorescence figures shown in figure legends, brightness and contrast were post-processed using the IrfanView software to improve the qualities of the images.

### 2.11. Semi-quantitative analysis of fluorescence intensities

Full lengths of the retinas were captured for analysis. Fluorescence intensities were measured using Fiji software. Areas of interest were determined in fluorescence using the default threshold, and integrated optical density (IOD) of the region of interest was measured. The area was normalized to the length of RPE (pixel) in corresponding images to correct for varying lengths of RPE in each image.

### 2.12. Statistical Analysis

All statistical analyses were carried out using Prism 8 software (GraphPad Software, Inc., La Jolla, CA) and Excel (Microsoft, Inc., Redmond, WA). All data sets were tested for normal distribution to decide on using parametric or non-parametric testing. For non-normally distributed data, Mann–Whitney U test was used, and for normally distributed data student t-test was used. All the statistical analyses were performed using two-tailed tests. The null hypothesis was that the two groups were not significantly different. Values are given as mean ±standard deviation, p<0.05 was considered statistically significant.

## 3. Results

### 3.1. Different ultrastructural findings in the RPE of albino and pigmented *Abca4^-/-^* mice without treatment

In pigmented mice it became obvious that most of the lipofuscin was fused with melanin to form MLF (Fig. 1 A). Some of the properties that are associated with these granules in *Abca4^-/-^* mice have been described previously [1]. However, under high power magnification (50 000 – 140 000- fold) we found unique thin membranes (3 – 4 nm thick) in the form of lamellae inside the non-melanin part of the MLF granule (Fig.1A). These TLMs are regularly present in MLF granules and the cytoplasm (Fig.2A) of RPE cells, and are associated with residual bodies (Fig. 1 B, C). TLMs have also been found in Bruch’s membrane (not shown) and are present in the endothelial cells (Fig. 1B, C) of the choriocapillaris and the capillary lumen (not shown), and often adhere to red blood cells when present in vessels, as demonstrated in the choriocapillaris of aged humans [21].

**Fig. 1.**
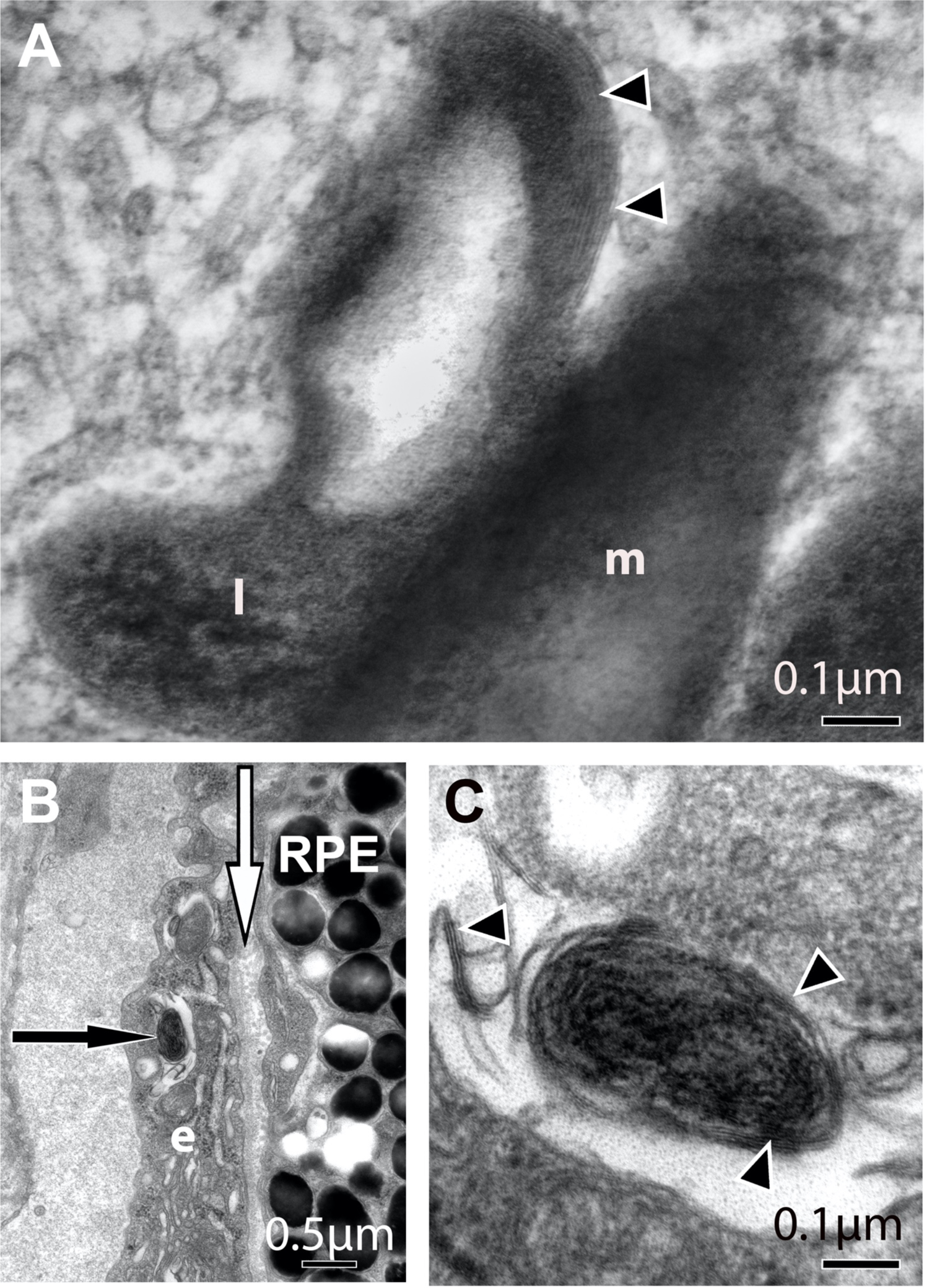
Detection of TLMs in MLF from pigmented *Abca4^-/-^* mice by high-resolution electron microscopy. A) A high power electron micrograph of an MLF granule from a pigmented *Abca4^-/-^* mouse magnified 50 000-fold is shown. The granule consists of melanin (m) with attached lipofuscin-like (l) material which is less electron-dense than the melanosome. Thin membranes form lamellae (arrowheads) in the upper part of this attachment. These thin membranes are regularly present in both the MLF granules and the cytoplasm of RPE cells and are associated with residual bodies (Fig. 1C). B): TLMs are found in Bruch’s membrane (white arrow) and are present in endothelial choriocapillaris cells (black arrow) and the capillary lumen (not shown). Fig.1 C shows the residual body within the endothelial cell in Fig 1 B surrounded by thin membranes (arrowheads) magnified 140 000-fold.

**Fig. 2.**
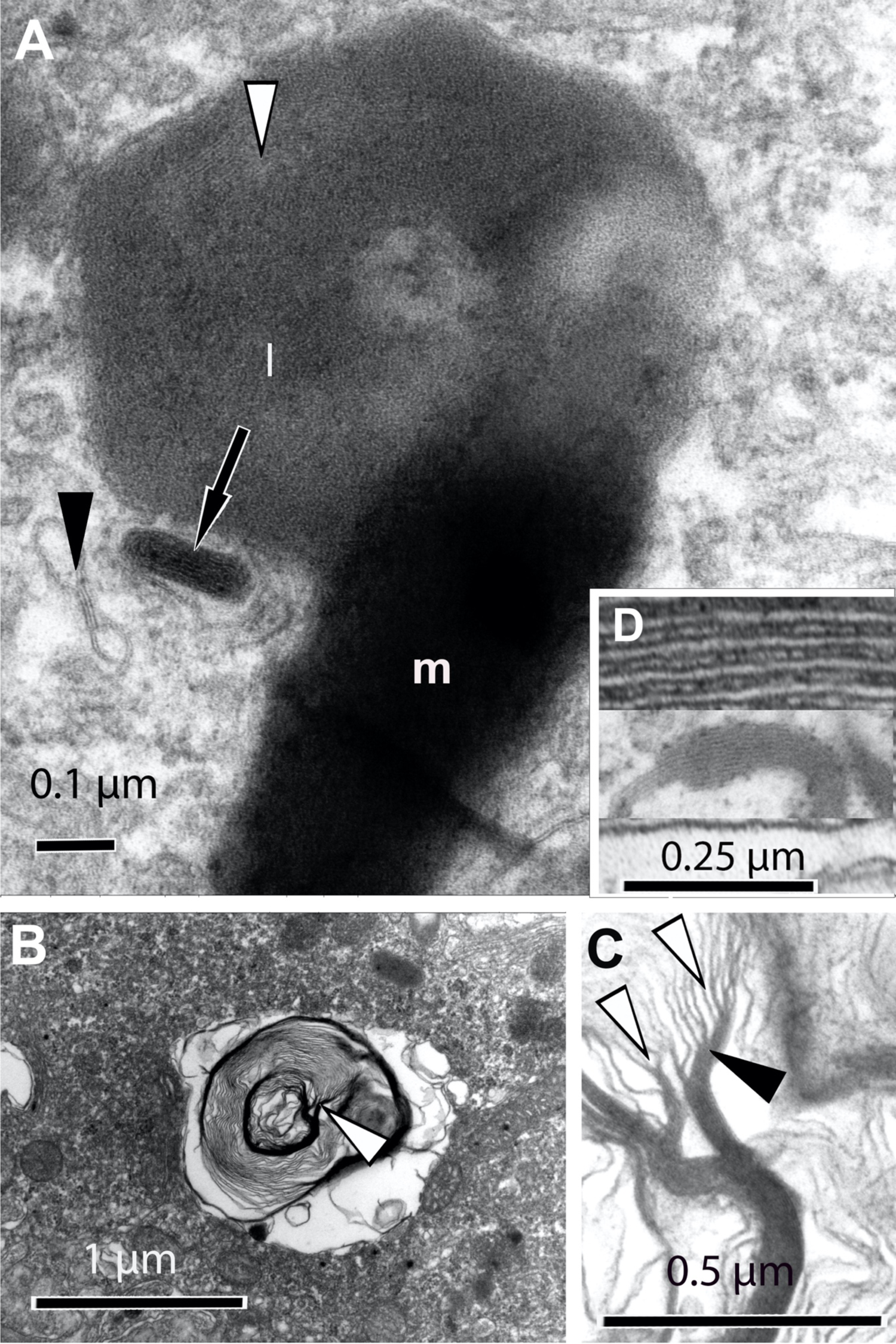
TLMs originate from photoreceptor disk membranes. A) A high power electron micrograph of a MLF granule from a pigmented *Abca4^-/-^* mouse magnified 50 000-fold. The granule consists of melanin (m) with attached lipofuscin-like (l) material which is less electron-dense than the melanosome. The lipofuscin content appears more homogeneous; however, fine lamellar membranes can be recognized in some areas (white arrowhead). The transition between melanin and lipofuscin is interwoven. A small fragment of the fine lamellar membrane is present within the cytoplasm (black arrowhead). Residual material which has the same appearance as the residual body in the endothelium (Fig.1 B, C) is associated with the MLF granules and surrounded with TLMs (arrow). It is not clear whether it is incorporated into or released from the MLF granule, but the latter is more likely because such structures are found in the endothelium and the bloodstream. Inset D) shows the proportion of the original photoreceptor disc membranes (upper part) and the condensed fine lamellar membranes under the same magnification (50 000-fold). B) A phagosome within the RPE cell of an albino *Abca4^-/-^* mouse is shown. Some of the phagosomal membranes are now electron-dense (arrowhead). C) Parts of the phagosome marked by the arrowhead in B. It is obvious that the original photoreceptor disk membranes (white arrowhead) have fused and become condensed (black arrowhead), and finally appear morphologically identical to the fine lamellar membranes in Fig.1,3.

MLF granules consisting of melanin and of lipofuscin-like material which is less electron-dense than the melanin, were observed. The lipofuscin content of MLF is generally homogeneous; however, the fine lamellar membranes (Fig.2A) were also observed. The transition between melanin and lipofuscin is interwoven (Fig.2A). The MLF granules resemble melanosomes after fusion with phagosomes, as seen in Fig.11 of an earlier report [9]. The original disc membranes inside the photoreceptors and phagosomes (Fig. 2B) fuse, and they become thinner and condensed (Fig. 2C) and resemble the TLMs found in the cytoplasm of RPE (Fig. 2 A) and endothelial cells (Fig.1 A, B).

The proportion of the original photoreceptor disc membranes and condensed fine lamellar membranes from a vacuole in the RPE of an albino *Abca4^-/-^* mouse are shown under the same magnification in Fig. 2D for comparison. Vacuole-like structures that have been observed in albino *Abca4^-/-^* mice are filled with heterogeneous types of materials (Fig.3A, B), with the majority composed of electron-dense or opaque TLMs (Fig.3C) and flocculate residues (Fig. 3A, B). These vacuole-like structures are separated from the cytoplasm by a membrane, which is generally incomplete or absent (Fig.3B). These vacuole-like structures are very frequently seen in albinos but are generally lacking in pigmented mice (Fig. 3D).

**Fig. 3.**
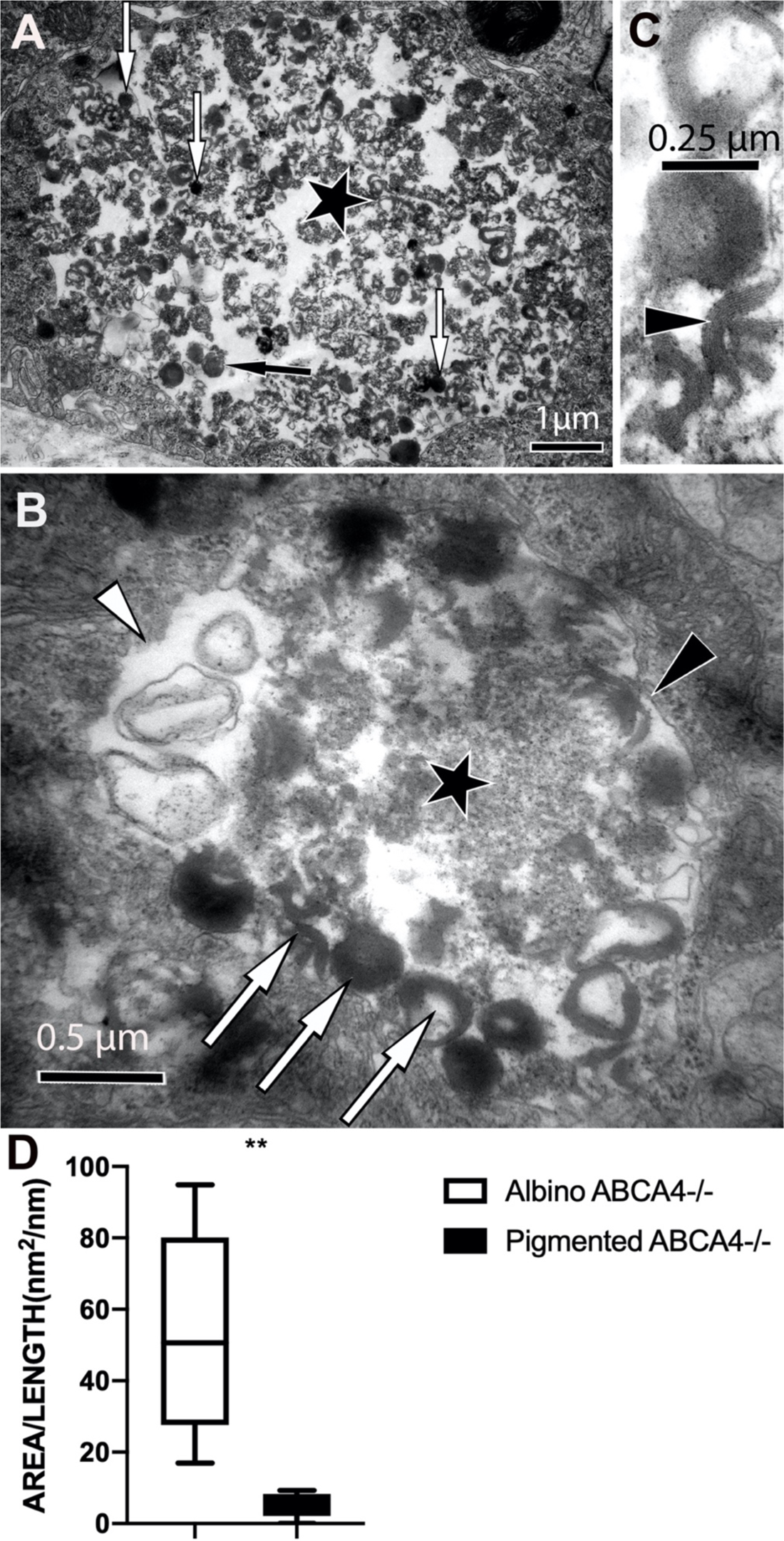
TLMs accumulate in the vacuoles of albino *Abca4^-/-^* mice. A, B) Vacuole-like structures (asterisks) that are often filled with heterogeneous material in albino *Abca4^-/-^* mice. The majority of the material inside these vacuoles consists of fine lamellar membranes (white arrows) that appear electron-dense (white arrow) or electron opaque (black arrow). Flocculate residues are also present (beside asterisk in B). These vacuole-like structures can be separated from the cytoplasm by membranes (black arrowhead); however, this membrane is often incomplete or absent (white arrowhead). C) The fine lamellar nature (arrowhead)(white arrow in B) of these residual bodies can be seen in the high-power micrograph. D): Quantification by electron microscopy revealed that these vacuole-like structures are barely present in pigmented *Abca4^-/-^*(7-month-old, *n* = 5 eyes/group, *p* < 0.01).

### 3.2. Electron microscopy of RPE cells from *Abca4^-/-^* mice following treatment with SIN-1

The intravitreal injection of SIN-1 was found to reduce the amount of lipofuscin present in the RPE cells of pigmented *Abca4^-/-^* mice (Fig.4 A-C), and melanin granules were more frequently in the form of single granules that were not associated with lipofuscin moieties (Fig.4B). The amount of melanin present was also reduced after SIN-1 treatment (Fig.4D). Many TLMs were detected at extracellular sites in the basal labyrinth following SIN-1 injection, indicating enhanced exocytosis (S1). Additionally, many small early-stage melanosomes appeared in the RPE of pigmented mice after SIN-1 treatment (S2). The reduction of lipofuscin-like material was not observed in the albino mice (Fig. 4 E-G). No evidence of toxicity was observed either, although this is not the focus of this study. Melanosomes were regularly observed within the lumen of the choriocapillaris (S3), indicating that melanin was removed from the RPE cells.

**Fig. 4.**
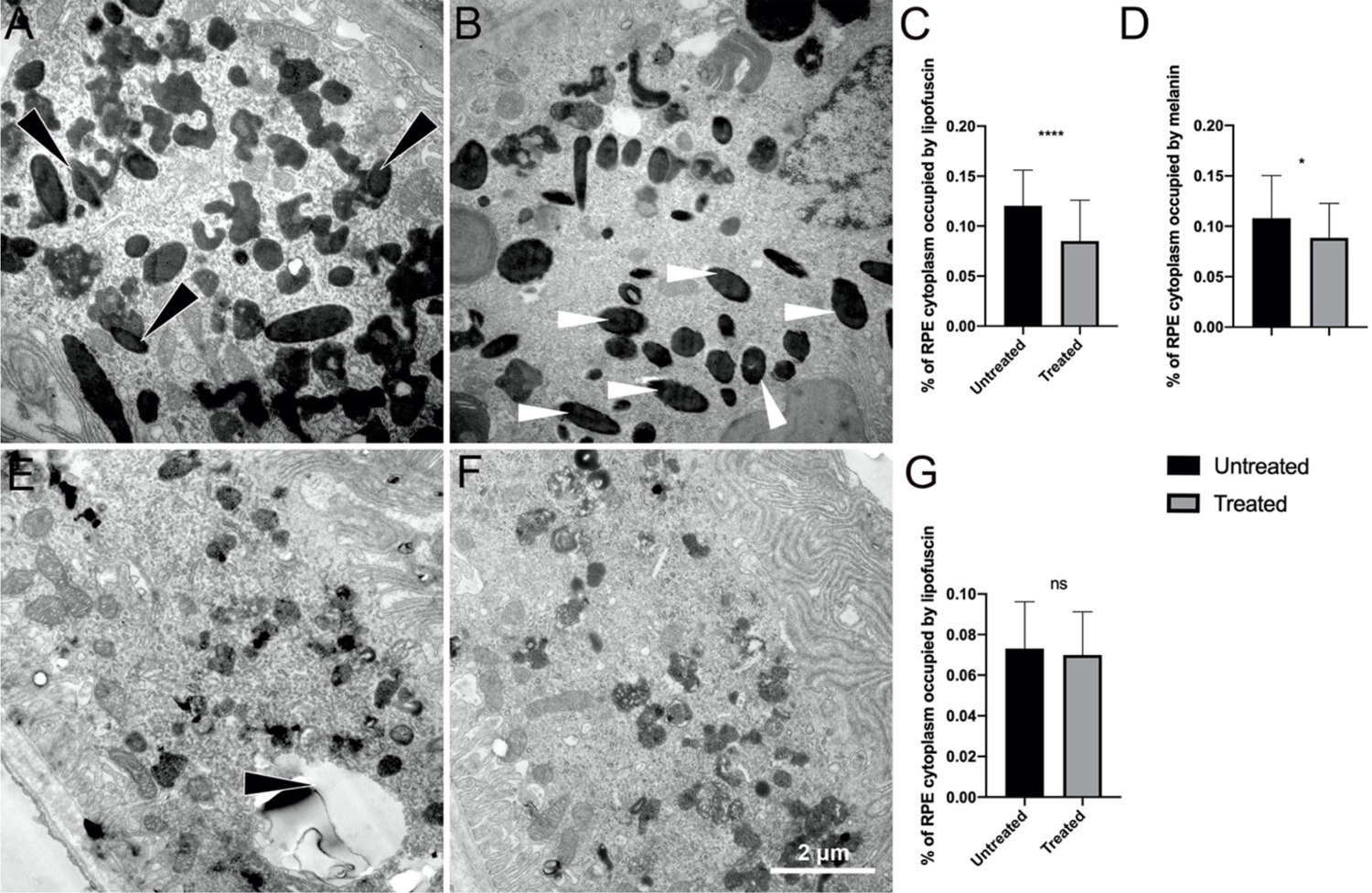
Lipofuscin is removed from the RPE of pigmented but not albino *Abca4^-/-^* mice after SIN-1 injection. A, B) Intravitreal injection of SIN-1 reduced the amount of lipofuscin in the RPE cells of pigmented *Abca4^-/-^* mice. A) Melanin granules fused with lipofuscin (black arrowheads). After intravitreal SIN-1 injection melanin granules occur more frequently as single granules that are not associated with lipofuscin moieties (white arrowheads in B) after treatment. A and B were taken from the eyes of the same mouse (B: right eye treated, A: left eye untreated). C) Quantification of the RPE area occupied by lipofuscin in untreated and treated pigmented *Abca4^-/-^* mice (*n* = 5 eyes/group, *p* < 0.0001). D) Quantification of the RPE area occupied by melanin in untreated and treated pigmented *Abca4^-/-^* mice(*n* = 5 eyes/group, *p* < 0.05). E-G) Reduction of the lipofuscin-like material was not observed in the albino mice. Fig. 4 E and F are taken from the eyes of the same mouse (E: left eye untreated, F: right eye treated). The black arrowhead in E marks a TLM in a vacuole. G) Quantification of the RPE area occupied by lipofuscin in untreated and treated albino *Abca4^-/-^* mice (*n* = 5 eyes/group, *p >0.05,* ns: not significant).

### 3.3. Electron microscopy of RPE cells from albino *Abca4^-/-^* mice after transfection with tyrosinase

Whereas the cytoplasm and vacuoles of the RPE cells in albino *Abca4^-/-^* mice were found to contain a substantial number of TLMs and large amounts of lipofuscin-like material (Fig.3, 5 A), this was reduced after melanin granules formed after transfection with tyrosinase (Fig.5 B). Typical premelanosomes indicating the melanogenic stages of choroidal melanogenesis were also formed in the choroidal melanocytes (S4).

**Fig. 5.**
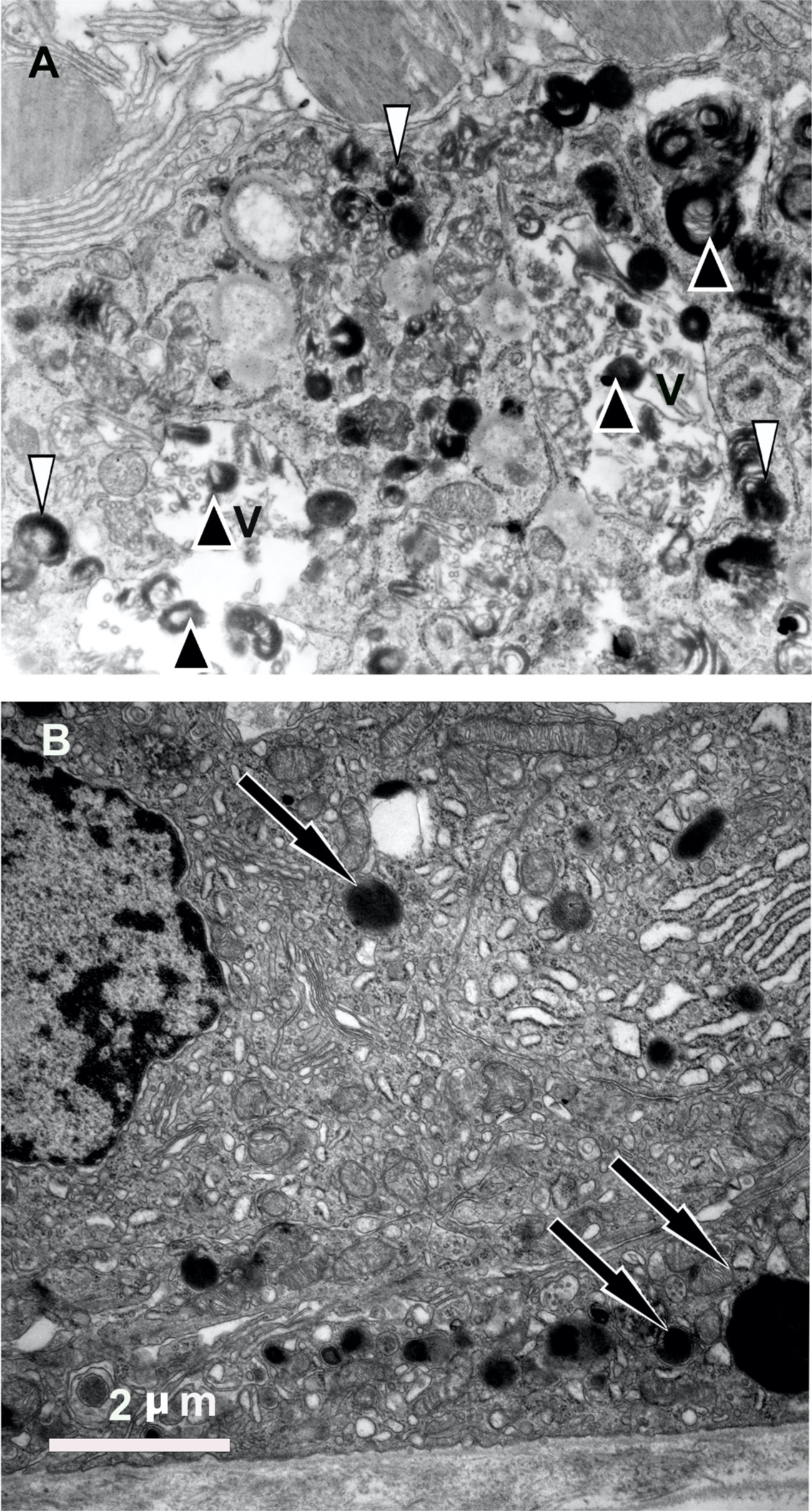
The amount of lipofuscin is reduced in the RPE cells that have formed melanin. Electron micrograph in A) shows typical lamellar TLMs (black arrowheads) that often accumulate in the vacuoles (V) and cytoplasm (white arrowheads) of RPE from albino *Abca4^-/-^* mice. B)The number of TLMs is reduced in the RPE of albino *Abca4^-/-^* mice that have been transfected with tyrosinase. The arrows in Fig.5 B point to newly formed melanosomes following transfection with tyrosinase.

### 3.4. Fluorescence microscopy and the fundus of albino *Abca4^-/-^* mice after transfection with tyrosinase

Melanosomes that were formed in the choroid and RPE of the albino *Abca4^-/-^* mice following transduction with tyrosinase via the subretinal injection of Ad- Tyr are shown in Fig.6A. Areas in which newly-formed melanosomes were present in the RPE and choroid produced NIR-AF signals, while no NIR-AF signals were generated in the absence of melanosomes (Fig.6A, S5). SW- AF shows that the lipofuscin signal almost disappeared in the area around the newly formed melanosomes as compared to areas with no pigmentation (Fig.6A). This effect is apparent in the magnified overlay of the NIR-AF and SW-AF images (Fig.6B).

**Fig. 6.**
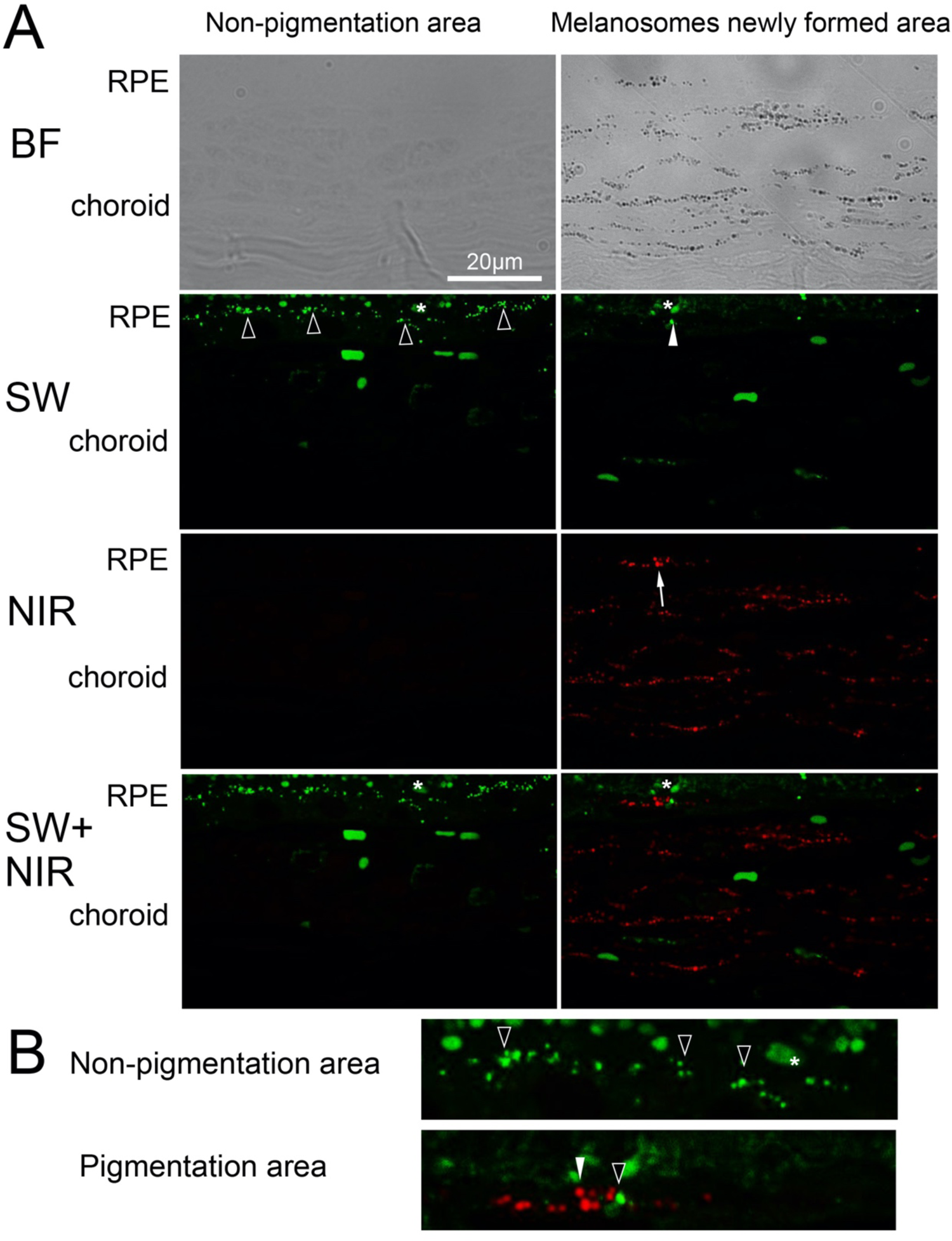
TLMs are reduced in albino Abca4-/- in RPE cells that have formed melanin. A) Overview of the RPE, choroidal BF, NIR-AF, and SW-AF. The BF image shows melanin granules that have formed after transfection with tyrosinase. No NIR-AF is observed in the non-pigmented area while newly formed melanosomes can be seen as NIR-AF in the pigmented area (white arrow). The number of lipofuscin granules is reduced in the pigmented area (white arrowhead) as compared to the non-pigmented area (black arrowhead). B) Magnified overlay of the NIR-AF and SW-AF images in the RPE areas shown in (A). The lipofuscin granules (black arrowhead) are almost invisible in the RPE cells with melanin formations (white arrowhead). Asterisks mark the autofluorescent outer segments. BF: bright field; NIR: near-infrared; SW: short wavelength.

### 3.5. Fluorescence microscopy of RPE from *Abca4^-/-^* mice after treatment with SIN-1, peroxidase, and ISDN

We found that the SW-AF intensity decreased in the eyes of pigmented *Abca4^-/-^* mice that were treated with SIN-1 as compared to untreated eyes (Fig. 7). Comparison of treated and untreated eyes from the same mice indicated that the lipofuscin levels are reduced after SIN-1 treatment, with semi-quantification revealing the same result (Fig.8A). A similar reduction in the SW-AF intensity was also observed after treatment with the other two reagents (Peroxidase, ISDN), which was performed in the same manner (Fig.8B, Fig.8C, S6). These results lead to the conclusion that the lipofuscin in pigmented *Abca4^-/-^* mice also can be degraded using NO generators and peroxidase.

**Fig. 7.**
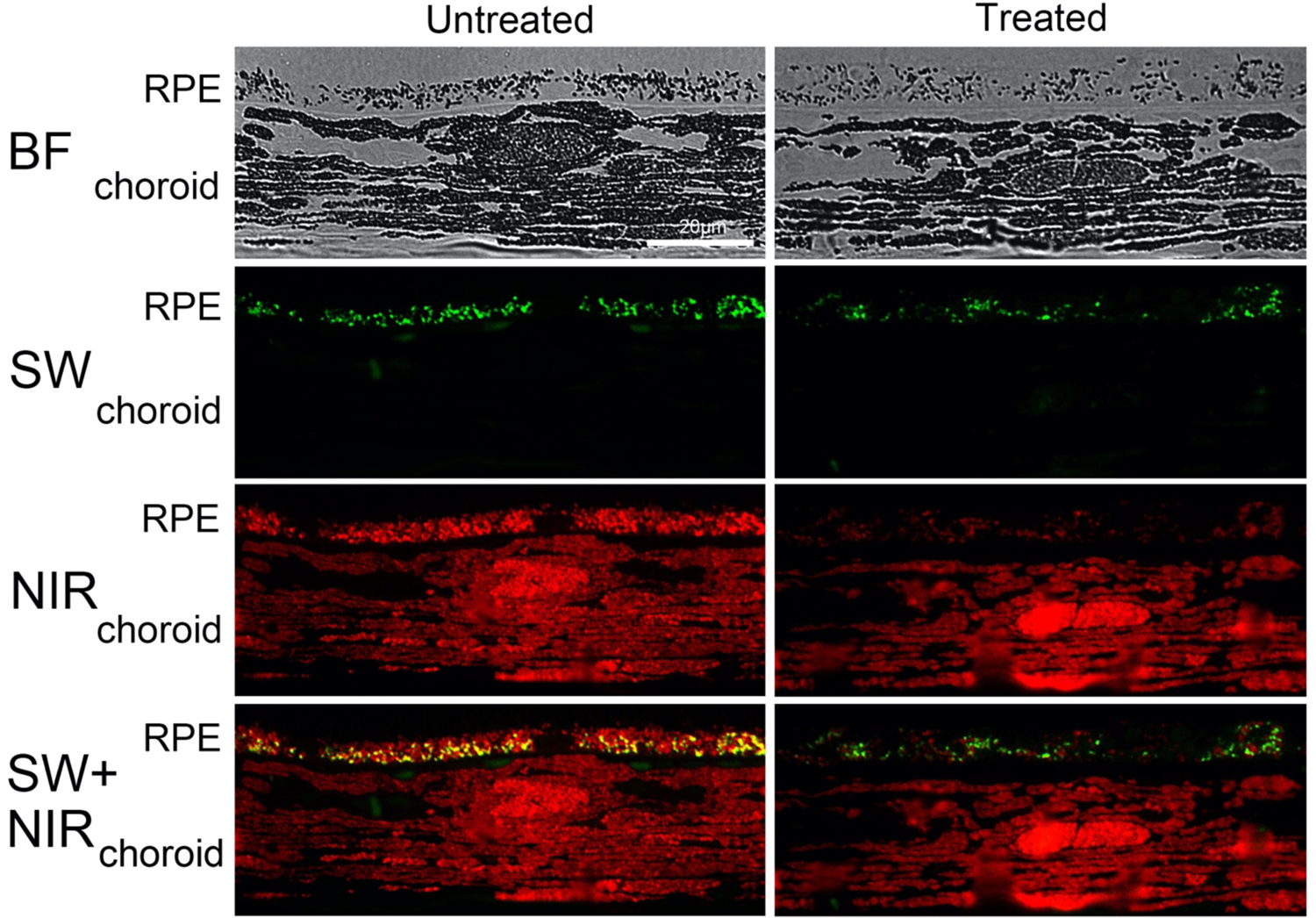
Lipofuscin is removed from RPE in pigmented *Abca4^-/-^* mice after SIN-1. A) After SIN-1 treatment, the amount of lipofuscin present is reduced. The yellow color shows areas where NIR-AF and SW-AF superimpose, representing MLF. The amount of MLF is also reduced after treatment. BF: bright field; NIR: near-infrared; SW: short wavelength.

**Fig. 8.**
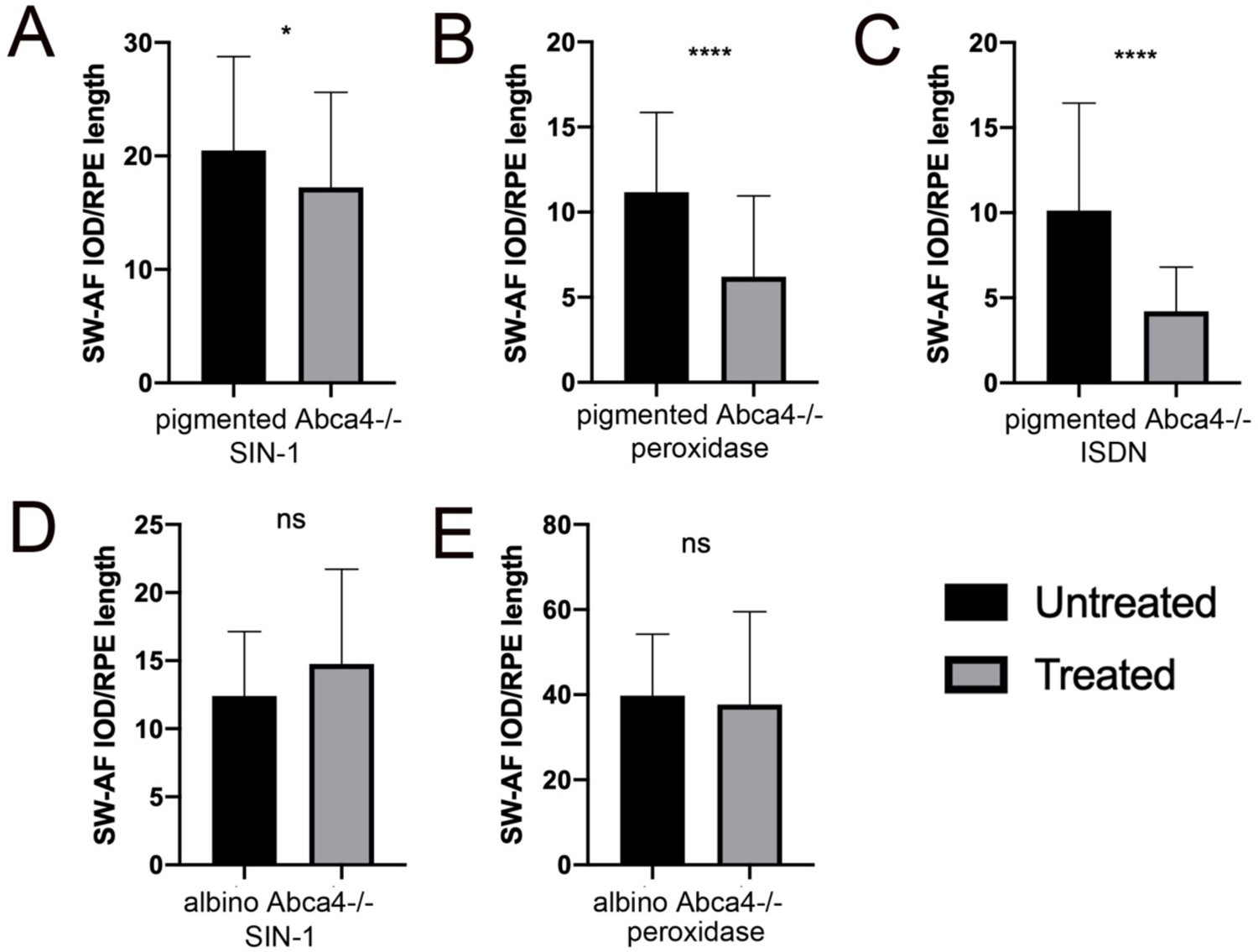
Quantification of lipofuscin. A) Semi-quantitative analysis of SW-AF intensities in RPE. The right eyes of the mice were treated with SIN-1(2 years old pigmented *Abca4^-/-^* mice, n=6 eyes/group, p<0.05). B) Semi-quantitative analysis of SW-AF intensities of RPE. The right eyes of the mice were treated with peroxidase (7-month-old pigmented *Abca4^-/-^* mice, n=3 eyes/group, P<0.0001). C) Semi-quantitative analysis of SW- AF intensities of RPE. The right eyes of the mice were treated with ISDN (7- month-old pigmented *Abca4^-/-^* mice, n=3 eyes/group, P<0.0001). D) Semi- quantitative analysis of SW-AF intensities of RPE. The right eyes of the mice were treated with SIN-1(6-month-old albino *Abca4^-/-^* mice, n=5 eyes/group, p>0.05, ns: not significant). E) Semi-quantitative analysis of SW-AF intensities of RPE. The right eyes of the mice were treated with peroxidase (7month-old albino *Abca4^-/-^* mice, n=3 eyes/group, p>0.05, ns: not significant).

**Fig. 9.**
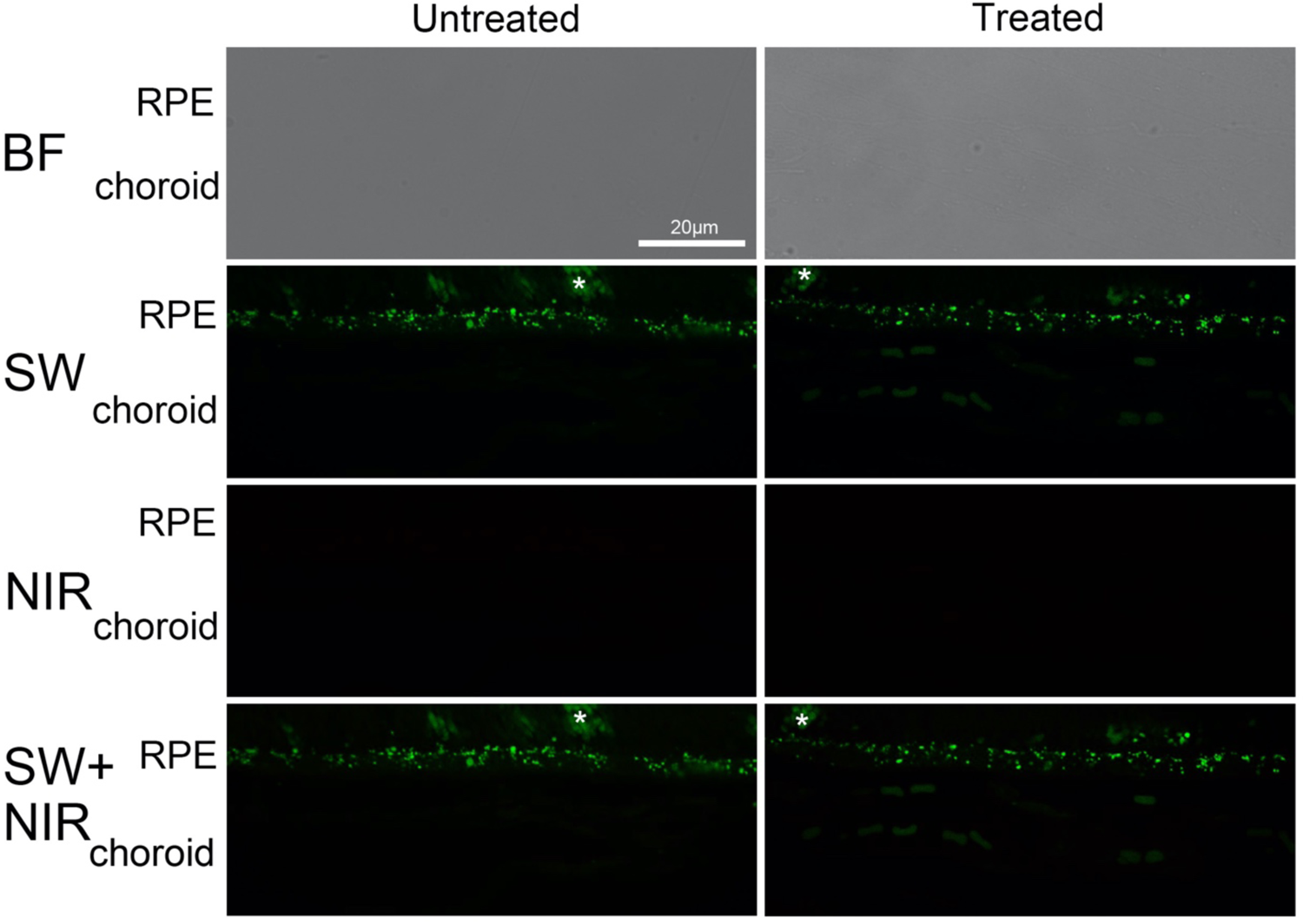
Lipofuscin is not removed from RPE of albino *Abca4^-/-^* mice after SIN-1. Reduction of lipofuscin granules was not observed in the albino *Abca4^-/-^* mice. No NIR-AF signal is observed due to the lack of melanin. Asterisks mark autofluorescent outer segments. BF: bright field; NIR: near-infrared; SW: short wavelength.

We next investigated whether melanin was required for lipofuscin degradation. To answer this question, the same treatment was applied to albino *Abca4^-/-^* mice. As shown in Figs.8D and 8E, neither SIN-1 nor peroxidase treatment affected the lipofuscin SW-AF intensity in albino mice..

## 4. Discussion

MLF formation starts with the formation of bisretinoids inside the disc membranes. The concentration of these bisretinoids is particularly high in animals and patients suffering from mutation in the Abca4 gene, which codes for a flippase that is localized in the disc membranes [22]. The excess retinal that forms in *Abca4^-/-^* mice reacts with phosphatidyletanol-amine inside the disc membranes to produce N-retinylidene-phosphatidylethanol-amine (N-ret-PE), which then reacts with all-trans-retinal to form bisretinoids. Such a reaction cannot be avoided and also takes place in young and healthy individuals. These molecules induce a variety of chemical reactions that end in the formatin of lipofuscin [23]. It is generally accepted that bisretinoids located within phagosomes cannot be digested by lysosomal enzymes. Nevertheless, the lack of lipofuscin formation in the RPE cells of young individuals indicates that some bisretinoid digestion may occur. This is also true for Stargardt patients, who can maintain normal vision for decades before their visual acuity decreases. This probably happens after the overload with lipofuscin has oxidized the melanosomes (see below).

Radicals are generated as part of the next steps in the formation of MLF. One of the most important functions of the RPE is the uptake and digestion of shed photoreceptor membranes that already contain bisretinoids, which is performed by phagosomes. Surprisingly, a large range of different radicals are formed during phagocytosis of the outer segments, with superoxide, hydroxyl radicals, hydrogen peroxide (H2O2), and the hazardous Fenton-type reaction occuring inside the phagosomes [24]. Radicals [25] can degrade bisretinoids [14, 26] and therefore some bisretinoid removal may take place during phagocytosis. However, phagosomal degradation is not able to digest all components completely particularly when there is an overload with bisretinoids as in *Abca4^-/-^* mice.

The fate of the incompletely degraded phagosomes was observed in the pigmented *Abca4^-/-^* mice models that were utilized in this study. The partially degraded phagosomes fuse with melanosomes, and the majority of the phagosomal membranes are converted into a homogenous material of different electron densities (Fig.1, 2) that accumulates within the melanolipofuscin granules. This observation has also been made in other species [8, 9, 27]. However, the digestion is not complete and some TLMs remain (Fig.1, 2), which are exocytosed and transported away with other residual materials via the choroidal blood vessels (Fig.1B, C, S3). These TLMs are very prominent in aged human eyes (in preparation), although only preliminary descriptions have been obtained [21]. They originate from the disc membranes in the photoreceptor cells, as demonstrated in Figs.2B, C.

The data obtained in this study indicate that the disc membranes continue to degrade within the melanosomes in the vicinity of melanin (Figs,1A,2A), where radical formation can also take place as outlined below. Melanosomes are specialized lysosomes that contain many lysosomal enzymes [28], which might indicate that their natural function is to degrade molecules. Eumelanin, commonly referred to as melanin, is a polymeric material that has captured the interest of various scientists for at least a century because of its unique physical and chemical properties. Neither these properties nor is its biological function are fully comprehended as yet. The complexity and properties of melanin indicate that it has significant potential as a functional polymeric material [29]. Melanin has unusual redox properties. It can both generate and absorb radicals and is a so-called stable radical [30, 31]. It can also change other types of energy into electric energy [32], and is known to be able to convert gamma radiation into chemical energy for growth and metabolism, as observed in the radiotrophic fungi detected in Chernobyl [33, 34]. Melanin is superoxide(O2-) donator [35] and serves as a naturally occurring biological source of electrons to power biochemical reactions [36].

Chemical interactions between lipofuscin and melanin have been repeatedly reported.

The chromophore bis-retinoid N-retinyl-N-retinylidene ethanolamine (A2E) [37], which is present in lipofuscin granules, is used as a model in research examining the properties of lipofuscin. The chemical composition of lipofuscin is currently unknown. A2E did not attenuate the antioxidant effects of DOPA-melanin and even potentiated it. The excess A2E in retinal pigmented epithelium cells was found to bind to the melanin in melanosomes, thus eliminating much of the toxicity [38]. The de novo synthesis of melanin in ARPE19 cells could be induced by feeding the cells with A2E and rod outer segments [39].

The concentration of paramagnetic centers in the MLF granules of human RPE is lower than that in melanosomes, which indicates that the melanin undergoes a destructive process in these granules. The irradiation of a mixture of melanosomes and lipofuscin granules with blue light (450 nm) results in the appearance of fluorescent melanin degradation products, which is in contrast to the results obtained when only melanosomes are irradiated. Dontsov et al. suggested that one of the main mechanisms of the age-related decrease in the melanin concentration of human RPE cells is its destruction in MLF granules under the action of superoxide radicals that form as a result of the photoinduced oxygen reduction that is performed by lipofuscin fluorophores [40].

The melanin that occurs in the RPE also contains Fe, Cu, and Zn [41] which all are involved in redox reactions. Fe has been shown to promote bisretinoid oxidation and the degradation of the bisretinoid A2E in the presence of light and reduce cell death in cell-based experiments. Moreover, the light-independent oxidation and degradation of A2E by Fenton chemistry products were evidenced by the consumption of A2E, release of dicarbonyls, and generation of oxidized A2E species in cell-free assays [42]. Lipofuscin and MLF form in the RPE of young rats who have been subjected to the removal of dietary zinc, and young individuals do not possess this type of granule under normal conditions [43]. Taken together, these reports present evidence for a relationship between the oxidizing capacity of melanin and lipofuscin degradation or formation. Another molecule that strongly interacts with melanin in the RPE is riboflavin, which is also a superoxide generator. Interestingly, no changes in riboflavin fluorescence have been detected after it binds with melanin. It appears that, contrary to cationic photosensitizing dyes, the singlet excited state of flavin molecules is not quenched by melanin, which might indicate a synergetic mode of action [44]. The oral substitution with riboflavin was found to remove lipofuscin from the RPE in pigmented *Abca4^-/-^* mice [12], indicating that riboflavin is involved in lipofuscin degradation. Riboflavin is known to play a role in the RPE because specific transporters for up taking riboflavin are present within the RPE, and reduced levels of these flavins result in the disruption of intracellular mechanisms, leading to photoreceptor cell death [45, 46] by a still unknown mechanism.

The drug Soraprazan which binds to melanin and melanolipofuscin has also been shown to form superoxides under light irradiation [20], which remove lipofuscin from the RPE of monkeys [11].

In addition, the results of this study show that autofluorescent lipofuscin could be removed from melanolipofuscin granules in pigmented *Abca4^-/-^* mice by treatment with NO generators or peroxidase. The finding that different NO donators could remove autofluorescent lipofuscin from the RPE of pigmented, but not albino, *Abca4^-/-^* mice is evidence for a mechanism that may be similar to chemiexcitation [16] and underlines the role of melanin. The mechanism appears to include the reaction of NO with a superoxide that is derived from melanin or lipofuscin to produce peroxynitrite, degrading the melanin to produce reactive compounds such as dioxetane that may be able to destroy bisretinoids. SIN-1 was found to moderately reduce the amount of melanin present in RPE cells (Fig.4D), which also indicates that melanin may have a role in reacting with NO. Whether lipofuscin degradation is a result of chemiexcitation or the generation of radicals induced by the reaction of NO with melanin remains unclear. But it is clear that melanin can generate superoxide which alone is able to degrade A2E-bisretinoid [13].

Nitric oxide is also formed naturally in the RPE of humans because NO synthetases are present in RPE cells [47] and nitro-A2E accumulates in the Bruch’s membrane of aged humans [48]. NO is also generated during the irradiation of melanin [49] and melanogenesis [50]. Tyrosinase, the key enzyme involved in melanogenesis, is up-regulated by nitric oxide [51]. Many reports have shown that the generation of NO is associated with both the enhancement [52] and the inhibition of melanogenesis [53, 54].

Radicals enhance the melanogenesis rate, which is the oxidation of L-tyrosine to dopachrome by tyrosinase, 40-fold, indicating a relationship between the presence of radicals and melanogenesis. All of these data support the view that NO can induce chemical reactions in MLF that degrade lipofuscin.

Peroxidase was found to very efficiently remove autofluorescent lipofuscin from the RPE of pigmented *Abca4^-/-^* mice in this study. Peroxidase can also eliminate A2E from RPE cells *in- vitro* [14]. Peroxidative activity has been observed in the melanosomes of RPE cells in cattle [8]. Therefore it is not unexpected for peroxidase to remove lipofuscin from melanolipfuscin granules in this study. Peroxidase may have substituted for the diminished peroxidative activity of melanin. The role of melanosomes in lipofuscin turnover also becomes evident if the findings for pigmented and albino *Abca4^-/-^* mice are compared. Albino *Abca4^-/-^* mice develop retinal pathology whereas pigmented mice do not, which points to a protective role for melanin during lipofuscin overload [1]. The number of TLMs is much higher in albino *Abca4^-/-^* mice as compared to pigmented *Abca4^-/-^* mice (Fig.3D), and the evidence indicates that TLMs can be digested to some extent inside melanolipofuscin granules. Whether NO generation also has an effect on the number of TLMs produced is currently unknown.

Moreover, the morphology of the lipofuscin granules in the two strains differs significantly [1] (Fig.4), indicating a role for melanin in the formation of lipofuscin. The lipofuscin part of the MLF granule is more homogeneously degraded to smaller units in the pigmented mice as compared to the lipofuscin residues in the albino mice. The lipofuscin components do not really form granules in the albinos, and are often missing a limiting membrane (see Fig.4b in [1]). Lipofuscin is also removed from the artificially pigmented RPE cells of albino *Abca4^-/-^* mice (Fig.6), and the newly formed melanosomes become oxidized and fluorescent in the NIR range (Fig. 6, S2).Furthermore, NO generators do not remove lipofuscin without the presence of melanin. The same is true for peroxidase.These data present further evidence for the existence of chemical reactions between melanin and lipofuscin.

The two strains differ in their genetic background: pigmented *Abca4^-/-^* mice are bred on a 129 background, while albino *Abca4^-/-^* mice have a BALB/c background. However, differences such as the number of TLMs, morphology of the lipofuscin, effects of NO generators, and the reaction to peroxidase are more likely caused by the melanin pigmentation than the genetic background of the strains [1] because our data show that melanin is associated with TLMs and lipofuscin.

Neither preliminary electroretinography (ERG) measurements nor ultrastructural histology provided evidence for the toxicity of the NO generators. We used ERG to investigate the effects that three NO generators (ISDN, Isosorbide mononitrate, and Pentaerythritol tetranitrate) had on the retinal function of *Abca4^-/-^* mice following intravitreal injection, but no evidence of toxicity was obtained [55]. It has also been reported that SIN-1 can protect the retinas of rats against toxicity when mediated by X-Ray irradiation [56]. However, investigating the potential toxicity of the NO generators is out of the scope of this study.

Several toxic products and radicals are formed when bisretinoid degradation occurs [13], meaning that it is advantageous for this process to occur in a closed compartment such as the melanosome in the vicinity of melanin because the toxic byproducts and radicals can be absorbed, thereby protecting the cell cytoplasm when such a reaction runs out of control. Nevertheless, melanin is affected by such reactions during aging and will itself become pro-oxidant [57], meaning that it loses the ability to remove lipofuscin and MLF begins to accumulate.

The visual acuity of Stargardt patients drops rapidly after the melanin granules become autofluorescent in the NIR mode [58]. This opens the door to speculate that that melanosomes have been oxidized by reactions with lipofuscin.

A similar mechanism has been observed in Abc*a4^-/-^* mice, which accumulate high numbers of MLF granules while the melanosomes become oxidized and fluorescent in the NIR range. This does not happen in age-matched wild-type mice [1], which also strongly supports the view that melanin is oxidized via a reaction with lipofuscin after overloading.

Here we show for the first time that melanin plays an important, if not the key, role in the degradation of fluorescent lipofuscin in RPE cells. The ability to remove autofluorescent lipofuscin from cultured human PRE cells by the superoxide generating drug Soraprazan was blocked by the radical scavenger Cardioxane [15] and lipofuscin removal was induced by NO generating drugs. Therefore, lipofuscin removal is more likely achieved by radical substitution than by enzymatic mechanisms.

## 5. Conclusions

Indigestable material resulting from the breakdown of photoreceptor membranes fuses with the melanosomes in the RPE [10] and forms MLF granules.

Autofluorescent lipofuscin is further eliminated from these granules by treatment with drugs that generate NO in pigmented, but not albino *Abca4^-/^*,*^-^* mice. These results suggest that melanin is involved in the degradation of lipofuscin.

## Supplementary material figure legends

**Extrusion of TLMs in the RPE of pigmented *Abca4^-/-^* mice after SIN-1 treatment**

**S1:** The arrowheads label TLMs that have extruded into the extracellular space of the basal labyrinth after treatment with SIN-1 in the RPE cells of pigmented *Abca4^-/-^* mice.

**SIN-1 induced melanogenesis in the RPE cells of pigmented *Abca4^-/-^* mice.**

**S2:** Many early-stage (black arrow) and more mature melanosomes (white arrow) are seen in the RPE. The inset shows the stages of melanogenesis at higher magnification

**Melanosomes are present within the choriocapillaris of pigmented *Abca4^-/-^*mice.**

**S3:** Several melanosomes (black arrowheads) are present within the lumen of choriocapillaris after treatment with SIN-1, indicating turnover of melanin in the RPE. This phenomenon could also be observed without SIN-1 treatment; however, it is likely that SIN-1 enhanced the rate of turnover. Also, electron opaque granules (black arrowheads) that resemble lipofuscin are localized within the blood vessel.

**Expression of tyrosinase-induced melanogenesis in choroidal melanocytes of albino *Abca4^-/-^* mice.**

**S4:** Many classical early-stage melanosomes are present in a choroidal melanocyte of an albino *Abca4^-/-^*^-^ mouse after transfection with the tyrosinase gene.

**A confocal scanning laser ophthalmoscope fundus AF image from an albino *Abca4^-/-^* mouse 2 weeks after injection with Ad-tyr**

**S5:** A) Fundus shows NIR-AF signals after Ad-tyr injection(right). The strong SW- AF signals are the result of tissue damage after injection (left). B) Without tyrosinase injection, NIR-AF is completely absent in albino *Abca4^-/-^* mice (Taubitz et al., 2018)(the same animal untreated eye). Ten out of twelve mice showed NIR-AF in the fundus.

**ISDN treatment removed lipofuscin from pigmented *Abca4^-/-^* mice**

**S6:** The amount of lipofuscin is reduced after treatment with ISDN. (A: untreated. B: treated.). The arrows mark lipofuscin granules in the RPE.

## Acknowledgments

The authors thank Antonina Burda for her technical assistance. This work was partially supported by the China Scholarship Council(CSC201906230325; to Yanan Lyu) and by the Deutsche Forschungsgemeinschaft BI 155/3-1 Projekt No. 401013432. The funders had no role in study design, data collection, analysis, decision to publish, or preparation of the manuscript. We also thank Douglas Brash for the helpful discussion.

## Abbreviations

Abca4: ATP-binding cassette A4
AMD: Age-related macular degeneration
A2E: N-retinyl-N-retinylidene ethanolamine
BF: Bright field
cSLO: confocal scanning laser ophthalmoscopy
ERG: Electroretinography
H_2_O_2_: Hydrogen peroxide
IOD: Integrated optical density
ISDN: sosorbide dinitrate
NIR-AF: Near-infrared autofluorescence
NIR-R: Near-infrared reflectance
NO: Nitric oxide
N-ret-PE: N-retinylidene-phosphatidylethanol-amine
MLF: Melanolipofuscin
OCT: Optical coherence tomography
OsO4: Osmium tetroxide
ROS: Rod outer segment
RPE: Retinal pigment epithelium
SIN-1: 3-morpholinosydnonimine
O_2_-: Superoxide
SW-AF: Short-wavelength autofluorescence
TLM: Thin lamellar membrane
UAC: Uranyl acetate

## Reference

1. Taubitz T, Tschulakow AV, Tikhonovich M, Illing B, Fang Y, Biesemeier A, et al. Ultrastructural alterations in the retinal pigment epithelium and photoreceptors of a Stargardt patient and three Stargardt mouse models: indication for the central role of RPE melanin in oxidative stress. PeerJ. 2018;6:e5215. doi:10.7717/peerj.5215

2. Taubitz T, Fang Y, Biesemeier A, Julien-Schraermeyer S, Schraermeyer U. Age, lipofuscin and melanin oxidation affect fundus near-infrared autofluorescence. EBioMedicine. 2019;48:592–604. doi:10.1016/j.ebiom.2019.09.048

3. Kim HJ, Montenegro D, Zhao J, Sparrow JR. Bisretinoids of the Retina: Photo-Oxidation, Iron-Catalyzed Oxidation, and Disease Consequences. Antioxidants (Basel). 2021;10(9):1382. doi:10.3390/antiox10091382

4. Kim HJ, Sparrow JR. Bisretinoid phospholipid and vitamin A aldehyde: shining a light. J Lipid Res. 2021;62:100042. doi:10.1194/jlr.TR120000742

5. Fang Y, Tschulakow A, Taubitz T, Illing B, Biesemeier A, Julien-Schraermeyer S, et al. Fundus autofluorescence, spectral-domain optical coherence tomography, and histology correlations in a Stargardt disease mouse model. FASEB J. 2020;34(3):3693–3714. doi:10.1096/fj.201901784RR

6. Feeney-Burns L Fau-Hilderbrand ES, Hilderbrand Es Fau-Eldridge S, Eldridge S. Aging human RPE: morphometric analysis of macular, equatorial, and peripheral cells. Invest Ophthalmol Vis Sci. 1984;25:195–200.

7. Warburton S, Davis WE, Southwick K, Xin H, Woolley AT, Burton GF, et al. Proteomic and phototoxic characterization of melanolipofuscin: Correlation to disease and model for its origin. Mol Vis. 2007;13:318–329..

8. Schraermeyer U, Stieve H. A newly discovered pathway of melanin formation in cultured retinal pigment epithelium of cattle. Cell Tissue Res. 1994;276(2):273–279. doi:10.1007/BF00306113

9. Schraermeyer U, Heimann K. Current understanding on the role of retinal pigment epithelium and its pigmentation. Pigment Cell Res. 1999;12(4):219–236. doi:10.1111/j.1600-0749.1999.tb00755.x

10. Schraermeyer U, Peters S, Thumann G, Kociok N, Heimann K. Melanin granules of retinal pigment epithelium are connected with the lysosomal degradation pathway. Exp Eye Res. 1999;68(2):237–245. doi:10.1006/exer.1998.0596

11. Julien S, Schraermeyer U. Lipofuscin can be eliminated from the retinal pigment epithelium of monkeys. Neurobiol Aging. 2012;33(10):2390–2397. doi:10.1016/j.neurobiolaging.2011.12.009

12. Schraermeyer U, Fang Y, Taubitz T, Tschulakow A. Degradation of lipofuscin in Stargardt mice can be enhanced by the superoxide generator riboflavin - a hypothesis for melanolipofuscin formation (abstract). Invest Ophthalmol Vis Sci. 2019;60(9).

13. Ueda K, Zhao J, Kim HJ, Sparrow JR. Photodegradation of retinal bisretinoids in mouse models and implications for macular degeneration. Proc Natl Acad Sci U S A. 2016;113(25):6904–6909. doi:10.1073/pnas.1524774113

14. Wu Y, Zhou J, Fishkin N, Rittmann BE, Sparrow JR. Enzymatic degradation of A2E, a retinal pigment epithelial lipofuscin bisretinoid. J Am Chem Soc. 2011;133(4):849–857. doi:10.1021/ja107195u

15. Schraermeyer, U.B., M; Senn-Bilfinger, J; Sturm, E; Hanauer; G, Methods for the determination of compounds or compositions for the treatment of lipofuscin related diseases and compounds or compositions, in *United States Patent No. 10*,495,630 B2. 2019, Katairo GMBH: Germany.

16. Premi S, Wallisch S, Mano CM, Weiner AB, Bacchiocchi A, Wakamatsu K, et al. Photochemistry. Chemiexcitation of melanin derivatives induces DNA photoproducts long after UV exposure. Science. 2015;347(6224):842–847. doi:10.1126/science.1256022

17. Hogg N, Darley-Usmar VM, Wilson MT, Moncada S. Production of hydroxyl radicals from the simultaneous generation of superoxide and nitric oxide. Biochem J. 1992;281 (Pt 2)(Pt 2):419–424. doi:10.1042/bj2810419

18. Blair HA. Dapagliflozin: A Review in Symptomatic Heart Failure with Reduced Ejection Fraction [published correction appears in Am J Cardiovasc Drugs. 2022 Jan;22(1):109]. Am J Cardiovasc Drugs. 2021;21(6):701–710. doi:10.1007/s40256-021-00503-8

19. Julien S, Kociok N, Kreppel F, Kopitz J, Kochanek S, Biesemeier A, et al. Tyrosinase biosynthesis and trafficking in adult human retinal pigment epithelial cells. Graefes Arch Clin Exp Ophthalmol. 2007;245(10):1495–1505. doi:10.1007/s00417-007-0543-3

20. Julien-Schraermeyer S, Illing B, Tschulakow A, Taubitz T, Guezguez J, Burnet M, et al. Penetration, distribution, and elimination of remofuscin/soraprazan in Stargardt mouse eyes following a single intravitreal injection using pharmacokinetics and transmission electron microscopic autoradiography: Implication for the local treatment of Stargardt’s disease and dry age-related macular degeneration. Pharmacol Res Perspect. 2020;8(6):e00683. doi:10.1002/prp2.683

21. Schraermeyer U, Addicks K, Kociok N, Esser P, Heimann K. Capillaries are present in Bruch’s membrane at the ora serrata in the human eye. Invest Ophthalmol Vis Sci. 1998;39(7):1076–1084.

22. Molday RS. ATP-binding cassette transporter ABCA4: molecular properties and role in vision and macular degeneration. J Bioenerg Biomembr. 2007;39(5-6):507–517. doi:10.1007/s10863-007-9118-6

23. Chen Y, Okano K, Maeda T, Chauhan V, Golczak M, Maeda A, et al. Mechanism of all-trans-retinal toxicity with implications for stargardt disease and age-related macular degeneration. J Biol Chem. 2012;287(7):5059–5069. doi:10.1074/jbc.M111.315432

24. Miceli MV, Liles MR, Newsome DA. Evaluation of oxidative processes in human pigment epithelial cells associated with retinal outer segment phagocytosis. Exp Cell Res. 1994;214(1):242–249. doi:10.1006/excr.1994.1254

25. Zhao J, Kim HJ, Ueda K, Zhang K, Montenegro D, Dunaief JL, et al. A vicious cycle of bisretinoid formation and oxidation relevant to recessive Stargardt disease. J Biol Chem. 2021;296:100259. doi:10.1016/j.jbc.2021.100259

26. Kim SR, Jockusch S, Itagaki Y, Turro NJ, Sparrow JR. Mechanisms involved in A2E oxidation. Exp Eye Res. 2008;86(6):975–982. doi:10.1016/j.exer.2008.03.016

27. Wavre-Shapton ST, Meschede IP, Seabra MC, Futter CE. Phagosome maturation during endosome interaction revealed by partial rhodopsin processing in retinal pigment epithelium. J Cell Sci. 2014;127(Pt 17):3852–3861. doi:10.1242/jcs.154757

28. Diment S, Eidelman M, Rodriguez GM, Orlow SJ. Lysosomal hydrolases are present in melanosomes and are elevated in melanizing cells. J Biol Chem. 1995;270(9):4213–4215. doi:10.1074/jbc.270.9.4213

29. Mostert AB. Melanin, the What, the Why and the How: An Introductory Review for Materials Scientists Interested in Flexible and Versatile Polymers. Polymers (Basel). 2021;13(10):1670. doi:10.3390/polym13101670

30. Mostert AB, Rienecker SB, Noble C, Hanson GR, Meredith P. The photoreactive free radical in eumelanin. Sci Adv. 2018;4(3):eaaq1293. doi:10.1126/sciadv.aaq1293

31. Sever RJ, Cope FW, Polis BD. Generation by visible light of labile free radicals in the melanin granules of the eye. Science. 1962;137(3524):128–129. doi:10.1126/science.137.3524.128

32. Hill HZ. The function of melanin or six blind people examine an elephant. Bioessays. 1992;14(1):49–56. doi:10.1002/bies.950140111

33. Dadachova E, Bryan RA, Huang X, Moadel T, Schweitzer AD, Aisen P, et al. Ionizing radiation changes the electronic properties of melanin and enhances the growth of melanized fungi. PLoS One. 2007;2(5):e457. doi:10.1371/journal.pone.0000457

34. Kothamasi D, Wannijn J, Van Hees M, Nauts R, Van Gompel A, Vanhoudt N, et al. Exposure to ionizing radiation affects the growth of ectomycorrhizal fungi and induces increased melanin production and increased capacities of reactive oxygen species scavenging enzymes. J Environ Radioact. 2019;197:16–22. doi:10.1016/j.jenvrad.2018.11.005

35. Lapina VA, Dontsov AE, Ostrovskiĭ MA. Generatsiia superoksida pri vzaimodeĭstvii melaninov s kislorodom [Generation of superoxides during the interaction of melanins with oxygen]. Biokhimiia. 1984;49(10):1712–1718.

36. Kim YJ, Wu W, Chun SE, Whitacre JF, Bettinger CJ. Biologically derived melanin electrodes in aqueous sodium-ion energy storage devices. Proc Natl Acad Sci U S A. 2013;110(52):20912–20917. doi:10.1073/pnas.1314345110

37. Eldred GE. Age pigment structure. Nature. 1993;364(6436):396. doi:10.1038/364396a0

38. Dontsov AE, Koromyslova AD, Sakina NL. Lipofuscin component A2E does not reduce antioxidant activity of DOPA-melanin. Bull Exp Biol Med. 2013;154(5):624–627. doi:10.1007/s10517-013-2015-6

39. Poliakov E, Strunnikova NV, Jiang JK, Martinez B, Parikh T, Lakkaraju A, et al. Multiple A2E treatments lead to melanization of rod outer segment-challenged ARPE-19 cells. Mol Vis. 2014;20:285–300.

40. Dontsov AE, Sakina NL, Ostrovsky MA. Loss of Melanin by Eye Retinal Pigment Epithelium Cells Is Associated with Its Oxidative Destruction in Melanolipofuscin Granules. Biochemistry (Mosc). 2017;82(8):916–924. doi:10.1134/S0006297917080065

41. Biesemeier A, Schraermeyer U, Eibl O. Chemical composition of melanosomes, lipofuscin and melanolipofuscin granules of human RPE tissues. Exp Eye Res. 2011;93(1):29–39. doi:10.1016/j.exer.2011.04.004

42. Ueda K, Kim HJ, Zhao J, Song Y, Dunaief JL, Sparrow JR. Iron promotes oxidative cell death caused by bisretinoids of retina. Proc Natl Acad Sci U S A. 2018;115(19):4963–4968. doi:10.1073/pnas.1722601115

43. Biesemeier A, Julien S, Kokkinou D, Schraermeyer U, Eibl O. A low zinc diet leads to loss of Zn in melanosomes of the RPE but not in melanosomes of the choroidal melanocytes. Metallomics. 2012;4(4):323–332. doi:10.1039/c2mt00187j

44. Kozik A, Korytowski W, Sarna T, Bloom AS. Interactions of flavins with melanin. Studies on equilibrium binding of riboflavin to dopa-melanin and some spectroscopic characteristics of flavin-melanin complex. Biophys Chem. 1990;38(1-2):39–48. doi:10.1016/0301-4622(90)80038-9

45. Kelley RA, Al-Ubaidi MR, Sinha T, Genc AM, Makia MS, Ikelle L, et al. Ablation of the riboflavin-binding protein retbindin reduces flavin levels and leads to progressive and dose-dependent degeneration of rods and cones. J Biol Chem. 2017;292(51):21023–21034. doi:10.1074/jbc.M117.785105

46. Said HM, Wang S, Ma TY. Mechanism of riboflavin uptake by cultured human retinal pigment epithelial ARPE-19 cells: possible regulation by an intracellular Ca2+-calmodulin-mediated pathway. J Physiol. 2005;566(Pt 2):369–377. doi:10.1113/jphysiol.2005.085811

47. Bhutto IA, Baba T, Merges C, McLeod DS, Lutty GA. Low nitric oxide synthases (NOSs) in eyes with age-related macular degeneration (AMD). Exp Eye Res. 2010;90(1):155–167. doi:10.1016/j.exer.2009.10.004

48. Murdaugh LS, Wang Z, Del Priore LV, Dillon J, Gaillard ER. Age-related accumulation of 3-nitrotyrosine and nitro-A2E in human Bruch’s membrane. Exp Eye Res. 2010;90(5):564–571. doi:10.1016/j.exer.2010.01.014

49. Yacout SM, McIlwain KL, Mirza SP, Gaillard ER. Characterization of Retinal Pigment Epithelial Melanin and Degraded Synthetic Melanin Using Mass Spectrometry and In Vitro Biochemical Diagnostics. Photochem Photobiol. 2019;95(1):183–191. doi:10.1111/php.12934

50. D’Acquisto F, Carnuccio R, d’Ischia M, Misuraca G. 5,6-Dihydroxyindole-2-carboxylic acid, a diffusible melanin precursor, is a potent stimulator of lipopolysaccharide-induced production of nitric oxide by J774 macrophages. Life Sci. 1995;57(26):PL401–PL406. doi:10.1016/0024-3205(95)02244-2

51. Sasaki M, Horikoshi T, Uchiwa H, Miyachi Y. Up-regulation of tyrosinase gene by nitric oxide in human melanocytes. Pigment Cell Res. 2000;13(4):248–252. doi:10.1034/j.1600-0749.2000.130406.x

52. Cals-Grierson MM, Ormerod AD. Nitric oxide function in the skin. Nitric Oxide. 2004;10(4):179–193. doi:10.1016/j.niox.2004.04.005

53. Park HY, Kosmadaki M, Yaar M, Gilchrest BA. Cellular mechanisms regulating human melanogenesis. Cell Mol Life Sci. 2009;66(9):1493–1506. doi:10.1007/s00018-009-8703-8

54. Sanzhaeva U, Vorontsova Y, Glazachev Y, Slepneva I. Dual effect of nitric oxide on phenoloxidase-mediated melanization. J Enzyme Inhib Med Chem. 2016;31(6):1063–1068. doi:10.3109/14756366.2015.1088843

55. Schraermeyer, U., Compound for the treatment and prophylaxis of a lipofuscin-associated disease and/or a disease associated with aged oxidized melanin, WIPO, Editor. 02.08.2021, Eberhardt Karls Universität Tübingen: Medizinische Fakultät: WO; PCT/EP2021/071553. p. 33.

56. Ozer MA, Polat N, Ozen S, Parlakpinar H, Ekici K, Polat A, et al. Effects of Molsidomine on Retinopathy and Oxidative Stress Induced by Radiotheraphy in Rat Eyes. Curr Eye Res. 2017;42(5):803–809. doi:10.1080/02713683.2016.1238943

57. Rozanowski B, Burke J, Sarna T, Rozanowska M. The pro-oxidant effects of interactions of ascorbate with photoexcited melanin fade away with aging of the retina. Photochem Photobiol. 2008;84(3):658-70. Photochem Photobiol. 2008;84(3):658–670. doi:10.1111/j.1751-1097.2007.00291.x

58. Kong X, West SK, Strauss RW, Munoz B, Cideciyan AV, Michaelides M, et al. Progression of Visual Acuity and Fundus Autofluorescence in Recent-Onset Stargardt Disease: ProgStar Study Report #4. Ophthalmol Retina. 2017;1(6):514–523. doi:10.1016/j.oret.2017.02.008

