## Supplementary figures and images for "Melanosomes degrade lipofuscin and precursors that are derived from photoreceptor membrane turnover in the retinal pigment epithelium—an explanation for the origin of the melanolipofuscin granule"

### Supplemental Figure S1

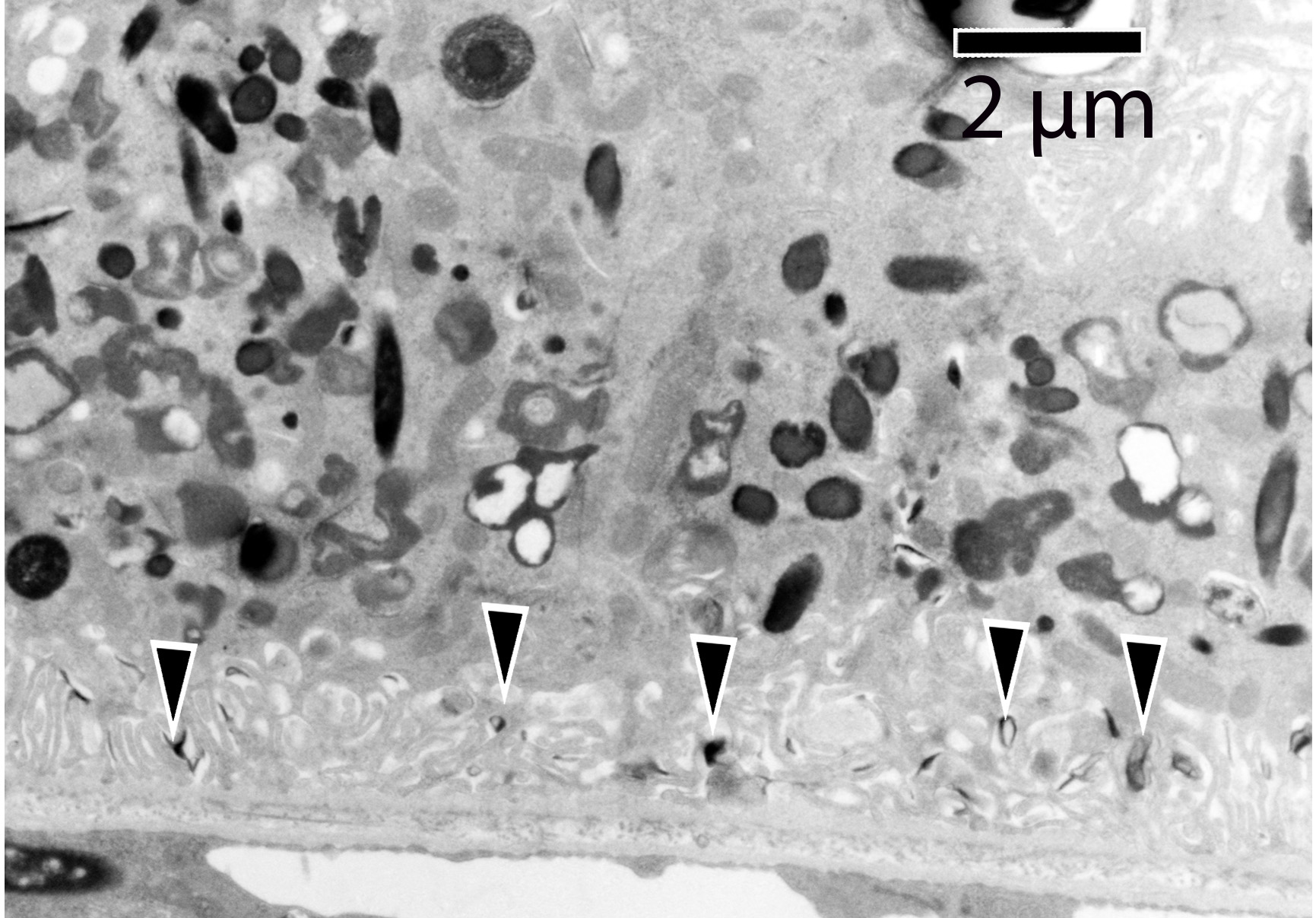

### Supplemental Figure S2

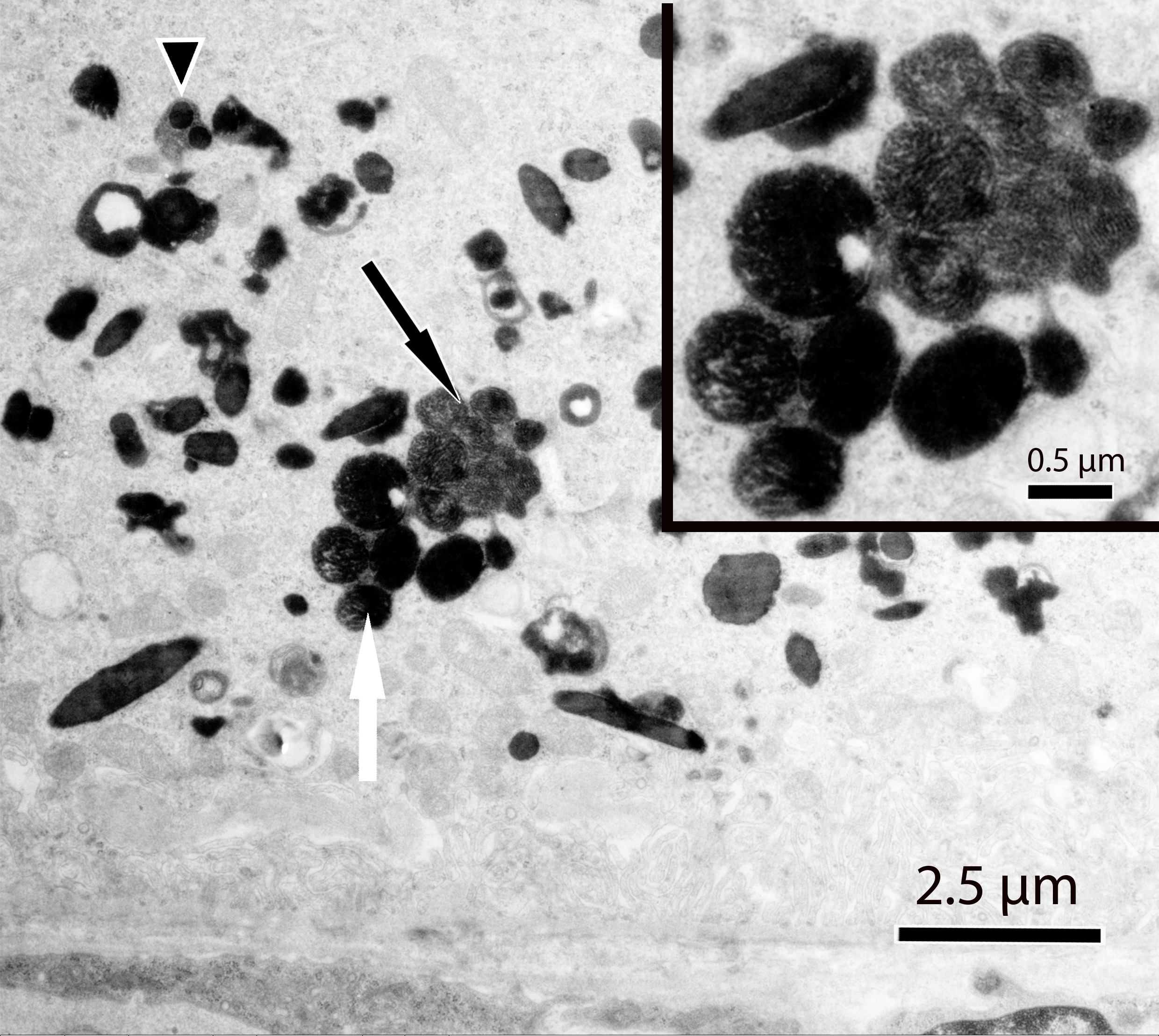

### Supplemental Figure S3

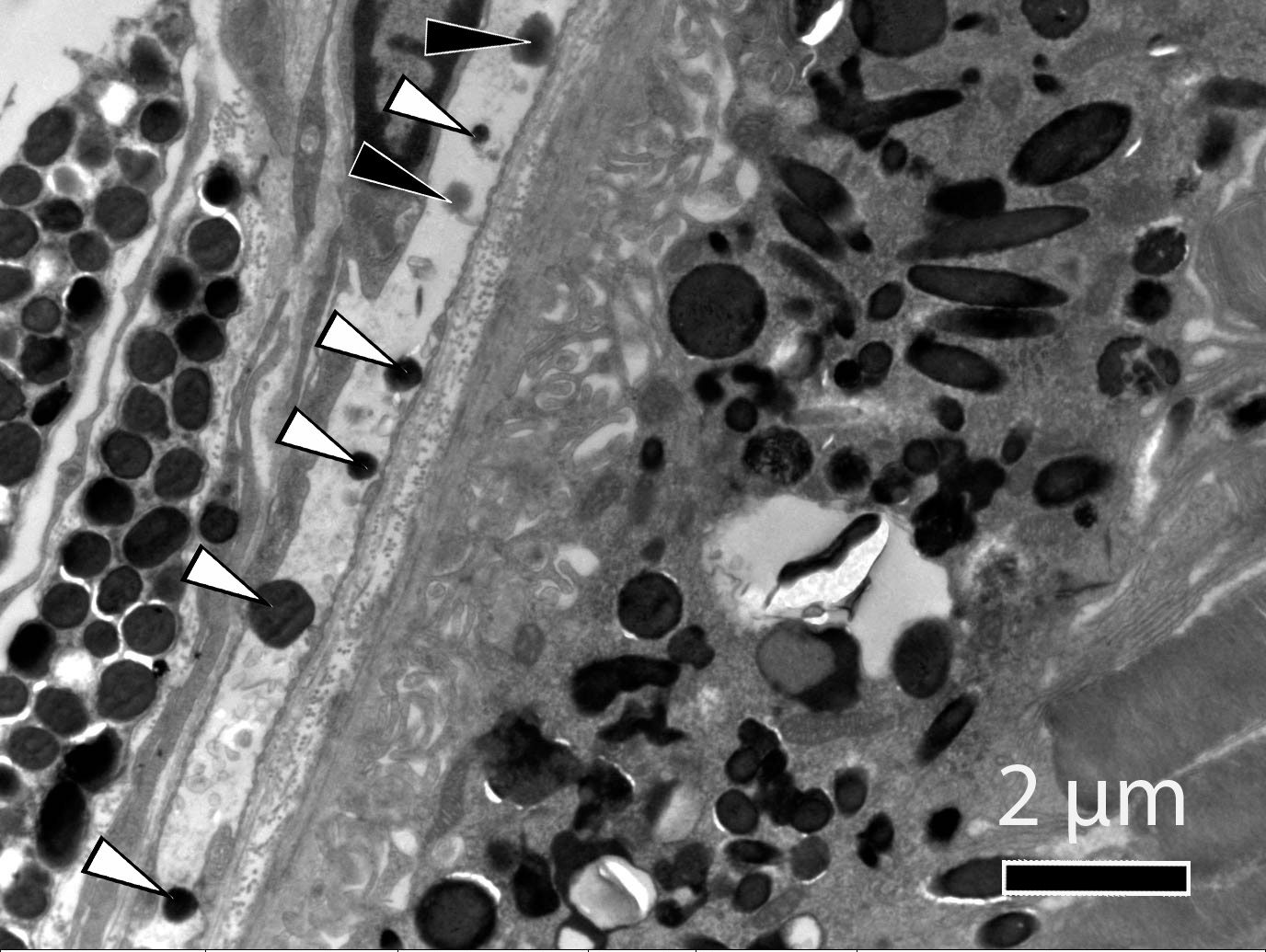

### Supplemental Figure S4

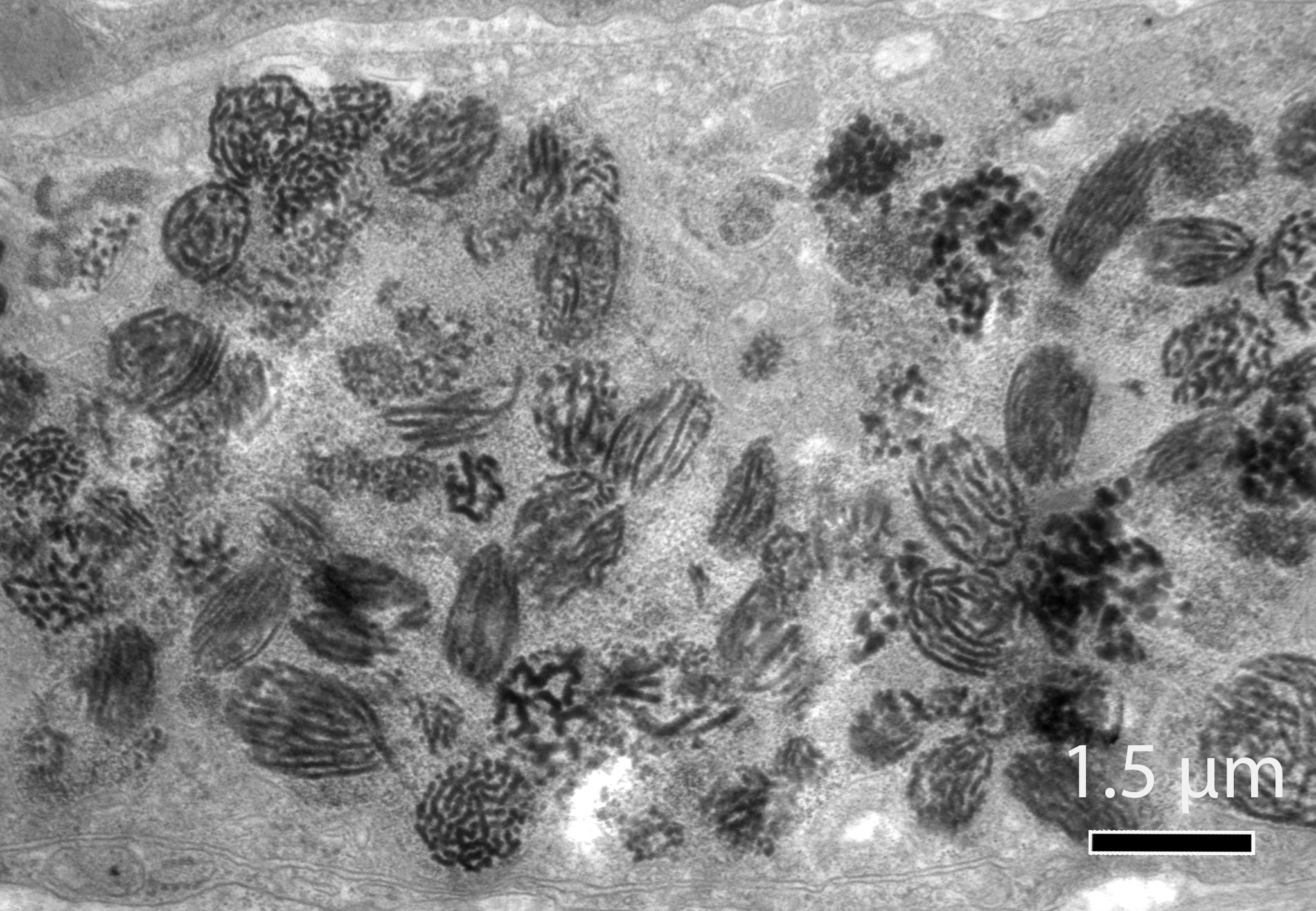

### Supplemental Figure S5

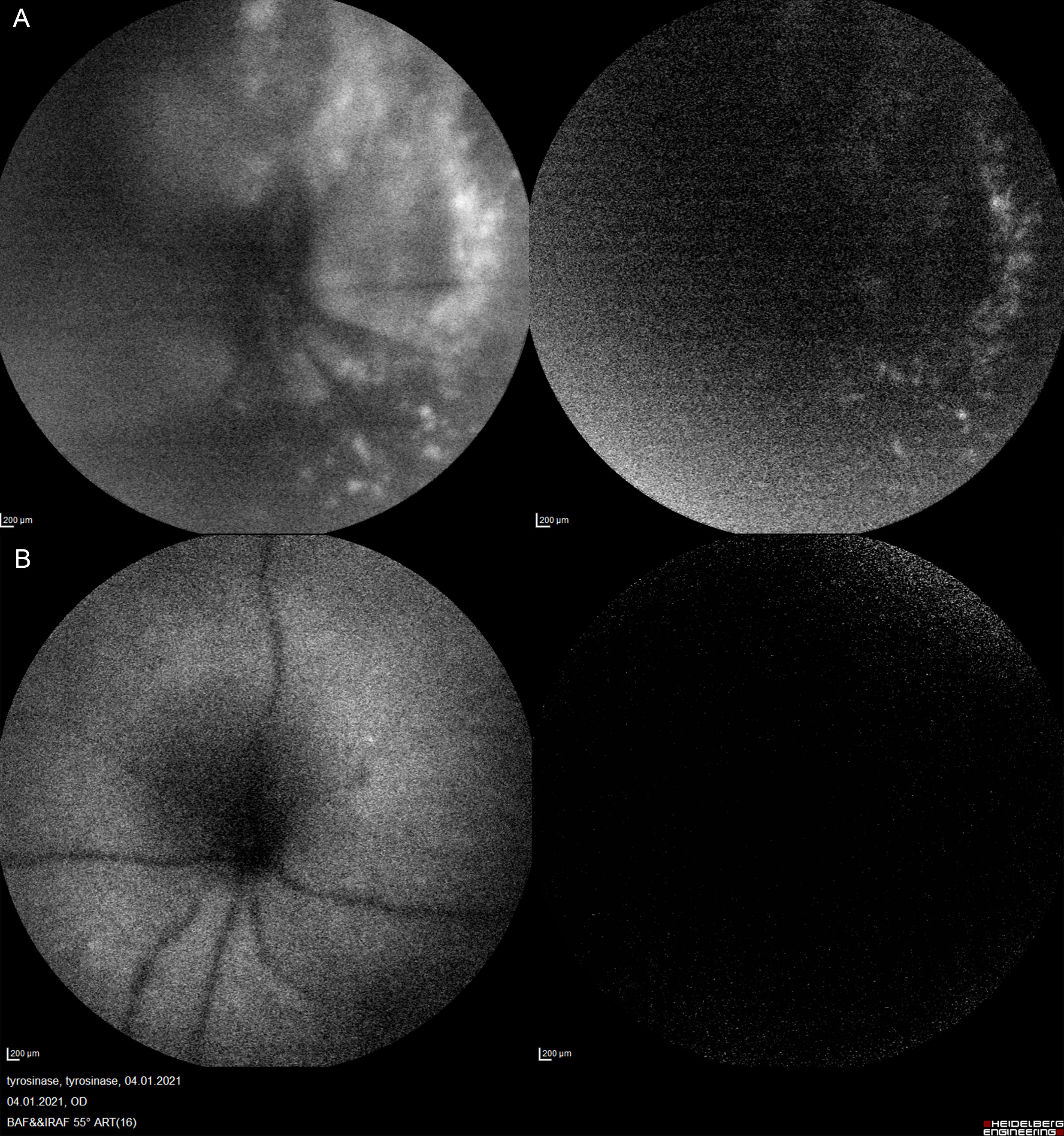

### Supplemental Figure S6

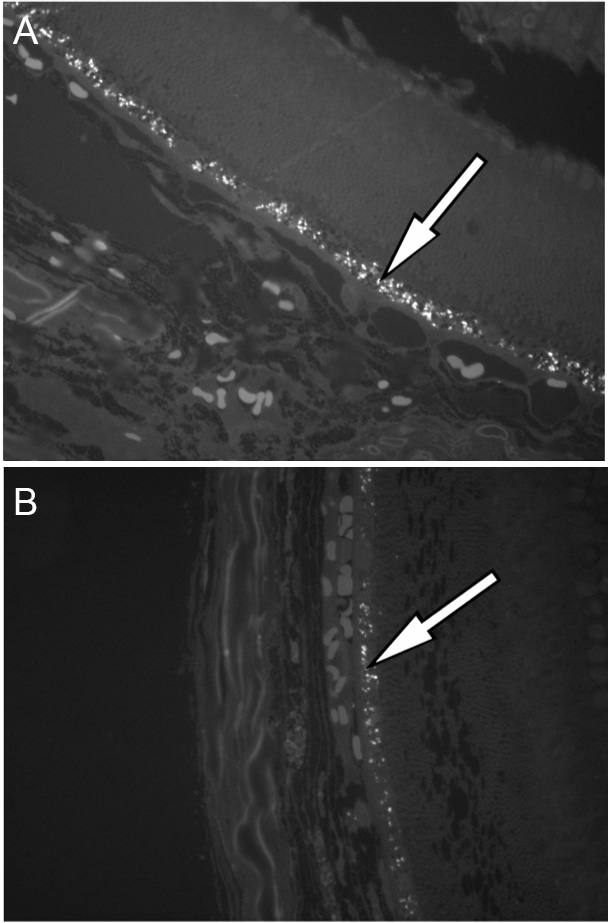
